# A novel toolkit for the efficient delivery of Cas9/sgRNA complexes to chromosomes in cells

**DOI:** 10.1101/812925

**Authors:** Ivana Indikova, Stanislav Indik

## Abstract

The application of gene editing technology is currently limited by the lack of safe and efficient methods to deliver RNA-guided endonucleases to target cells. We engineered lentivirus-based nanoparticles to copackage the U6-sgRNA template and the CRISPR-associated protein 9 (Cas9) fused with a virion-targeted protein Vpr (Vpr.Prot.Cas9), for simultaneous delivery to cells. In addition to Vpr, equal spatiotemporal control of the *vpr.prot.cas9* and *gag/pol* gene expression of was required for efficient packaging of the fusion protein into virus particles. Transduction of the unconcentrated, Vpr.Prot.Cas9-containing vectors resulted in >98% disruption of the *EGFP* gene in reporter HEK293-EGFP cells with minimal cytotoxicity. Furthermore, we detected indels in the targeted endogenous loci at frequencies of up to 100% in cell lines derived from lymphocytes and monocytes and up to 15% in primary CD4+ T cells by high-throughput sequencing. This approach may provide a platform for the efficient, safe and selective delivery of genome editing enzymes to cells and it may be suitable for simultaneous gene disruption and transgene delivery.

## Text body

The scope and scalability of gene editing systems is currently limited by problems with the delivery of the CRISPR/Cas RNA-guided endonuclease (RGEN) components to recipient cells. Lipofection, electroporation, nucleofection and virus-based techniques are widely used to deliver Cas9/sgRNA expression cassettes. Unfortunately, the methods for delivering DNA have limited cell-type specificity and are associated with side effects, such as integration into undesired chromosomal locations, immunogenicity, size-constrained packaging of expression cassettes (payload limit for AAV ≤4.7 kb) and increased off-targeting resulting from sustained expression. Increasing attention is thus being paid to the direct delivery of preassembled Cas9 protein/sgRNA complexes (RNPs) to cells ^1,2^, in which the rapid turnover of RNPs limits the exposure of the genome to nucleases, thereby mitigating off-target effects. Furthermore, the transient occurrence of RNPs in cells is expected to elicit minimal innate and adaptive immune responses, especially when a synthetic sgRNA lacking 5’ triphosphates and a Cas9 orthologue derived from a species other than *Streptococcus pyogenes* are used ^3,4^. Despite the advantages of RNP delivery, the use of this approach is restricted to cell types that do not suffer from reduced cell viability or phenotypic changes following chemical transfection or electroporation. Furthermore, the technology requires laborious optimization of the transfection protocol for every cell type and lacks tissue and cell specificity. Thus, there is an urgent need for a more versatile, safe, cell-selective and “easy-to-use” delivery system.

As a result of their efficiency, low toxicity, simplicity of production, mild immunogenicity, relative safety and ease of use and because of the possibility of customizing cell tropism, lentiviral vectors (LVs) are widely used in basic research and are being tested in numerous clinical trials for use in gene therapy (http://www.abedia.com/wiley/index.html) ^5,6^. Furthermore, LVs have recently been approved by the FDA for a genetic engineering of T lymphocytes for cancer immunotherapy ^7^. In addition to nucleic acids, LVs can also deliver foreign proteins of interest (POIs) to mammalian cells (reviewed in ^8^), and proof-of-concept studies have shown that LVs can serve as platforms for the administration of “protein-based” designer nucleases to ablate host genes ^9,10^. However, the system suffers from modest effectiveness and is not yet able to deliver the programmable nuclease and the sgRNA simultaneously. We now report an engineered LV that delivers the Cas9 protein and a template for sgRNA in “all-in-one transducing nanoparticles” and efficiently edits the targeted loci in the genome.

We postulated that that the large size (~160 kDa) and the net positive charge of Cas9 may cause structural disturbances and lead to reduced transducibility in such a case that the endonuclease is directly linked to the structural components of HIV-1 virions (embedded in the Gag polyprotein). To circumvent this problem, we translationally fused Cas9 containing an N-terminal protease cleavage site (Prot) to the C-terminus of an accessory protein of HIV-1, Vpr. Vpr interacts with the p6 domain of the Gag precursor, thereby mediating the encapsidation of its fusion partners into virions ^11^. Previous work with programmable nucleases such as meganucleases, which were packaged into lentiviral particles with the help of Vpr, showed only a moderate efficiency in genome editing, because little nuclease was packaged into virions ^9^. Earlier reports showed that retroviral mRNA nuclear export elements regulate protein function and virion assembly ^12–15^. Thus, we anticipated that it should be possible to increase the packaging of foreign proteins fused to Vpr, including Vpr.Prot.Cas9, by directing the transport dynamics and/or spatial distribution of the *vpr.prot.cas9* transcripts in the same manner as that of the *gag* mRNA. The matching spatiotemporal control of gene expression should localize the transcripts to the same cytoplasmic microdomain and result in co-localization of the nascent Vpr.Prot.Cas9 and Gag proteins. A close proximity of the proteins should promote their interaction and lead to enhanced encapsidation of the fusion protein to virions. We thus constructed two Vpr.Prot.Cas9 expression constructs carrying various RNA export functions, pVpr.Prot.Cas9 (containing the Rev responsive element, RRE) and pVpr.Prot.Cas9-CTE_3x_ (containing the constitutive transport element from MPMV, CTE; three copies of CTE were used to produce approximately the same levels of protein in the cytoplasm ^16^) (**Supplementary Figure 1a**). In the presence of Rev, the RRE facilitates mRNA export via the CRM1 pathway, whereas the CTE drives mRNA nuclear exit via the NXF1 pathway ^17–20^. The constructs were co-transfected into HEK293T cells together with four complementary plasmids: i) pHCMV-G, which produces the VSV.G envelope protein for pseudotyping virus particles; ii) psPAX2, a second-generation packaging construct, which provides the virion proteins; iii) pRSV-Rev, encoding Rev; and iv) the pLenti(sgRNA) transfer vector containing a U6 promoter driving the expression of an sgRNA specific to the targeted site (**Fig. 1a**). The amounts of Vpr.Prot.Cas9 produced in transfected cells and packaged into produced virions were determined by immunoblotting. The levels of the fusion protein in the cell lysates of cells transfected with the two constructs were the same but the amount of Vpr.Prot.Cas9 packaged into virions was markedly increased when the transcript contained the RRE (**Supplementary Fig. 1c**). The results support the view that the control of spatial localization and/or temporal distribution of transcripts encoding heterologous proteins may be used to facilitate packaging of the protein to virions (**Supplementary Fig. 1b**).

**Fig. 1:**
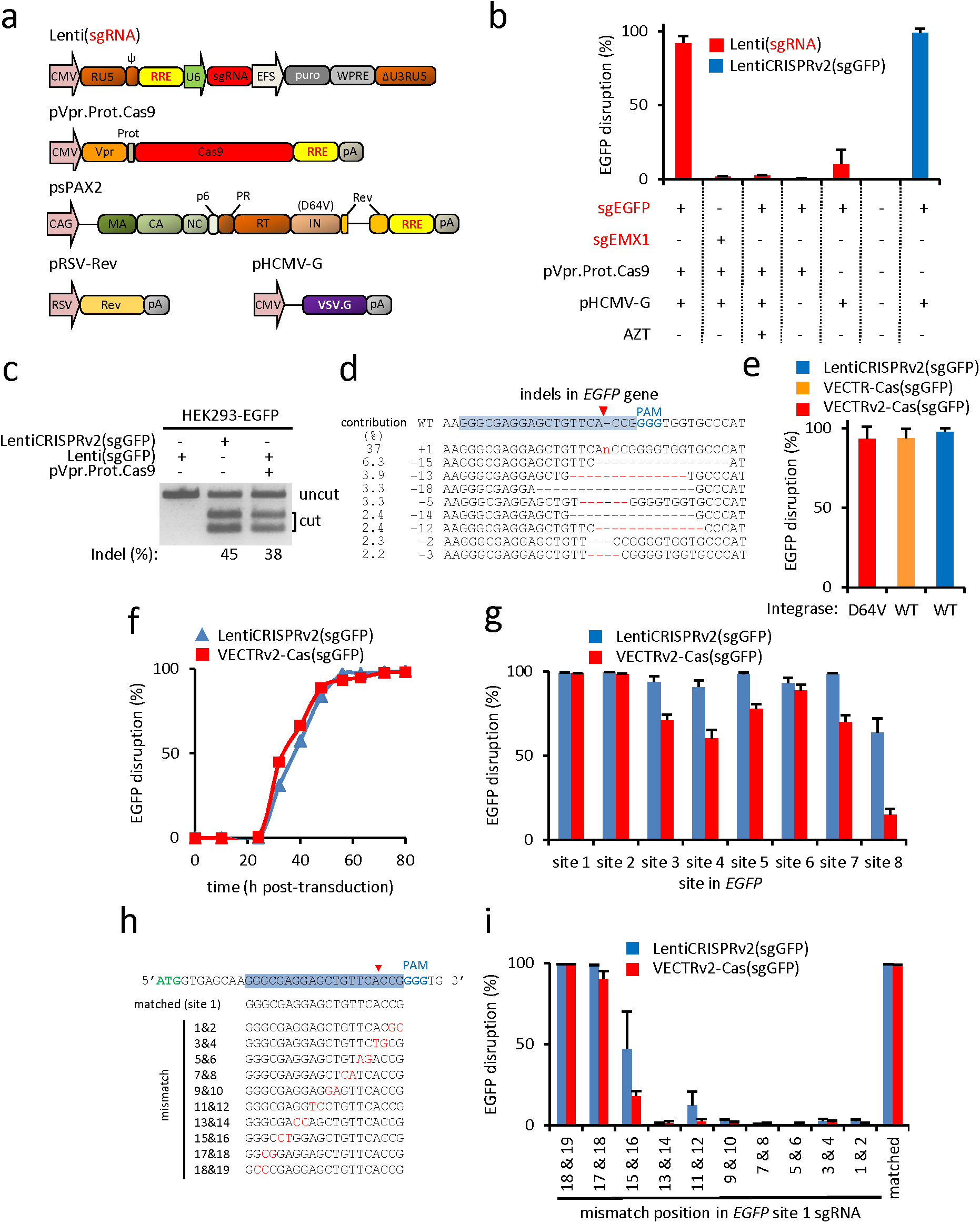
Delivery of two-component lentivector nanoparticles carrying the Cas9 nuclease protein and a template for the U6-sgRNA expression cassette to human HEK293-EGFP cells. **a,** Design of constructs to generate the lentivector particles. Cas9 was fused to the C-terminus of Vpr containing an authentic HIV-1 protease cleavage site (CTLNF/PISPI; Vpr.Prot.Cas9). The U6-sgRNA expression cassette was incorporated into a lentiviral expression vector (Lenti(sgRNA)). The packaging construct (psPAX2) encoded either wild-type or inactivated integrase (IN; D64V). The VSV.G envelope protein was used to pseudotype and stabilize viral particles (pHCMV-G). Efficient nuclear export and colocalization of mRNA for translation were supported by adding Rev-responsive element (RRE) to the constructs and by overexpressing Rev during virion production (pRSV-Rev). Gag-Pol subunits: matrix (MA), capsid (CA), nucleocapsid (NC), p6, reverse transcriptase (RT), and integrase (IN). Packaging signal (ψ); promoters (CMV, CAG, RSV, U6, and EFS), polyadenylation signal (pA), posttranscriptional regulatory element (WPRE). **b,** *EGFP* gene disruption in HEK293-EGFP cells after transduction with the two-component lentivectors (red entries) or a control LentiCRISPRv2(sgGFP) (blue entry). **c,** T7 endonuclease I (T7EI) assay to measure indels in the *EGFP* gene resulting from transduction with the two-component lentivector, the same vector lacking Vpr.Prot.Cas9 or the control pLentiCRISPRv2(sgGFP). **d,** Mutant sequences at the *EGFP* locus and their frequencies, as determined by SYNTHEGO analysis of Sanger sequencing of a PCR product amplified from VECTR-Cas(sgGFP)-transduced HEK293-EGFP cells. The 20-nt target sequence is shown with a blue background. The protospacer adjacent motif (PAM) sequence is shown in blue. **e,** Comparison of EGFP disruption after transduction with lentiviral particles containing integration-deficient (D64V) or integration-proficient (WT) integrase. **f**, Time course analysis of EGFP disruption mediated by the two-component VECTR-Cas(sgGFP) or the gene-delivering LentiCRISPRv2(sgGFP). **g**, EGFP disruption activity of the two lentivectors directed to different sites within the *EGFP* gene. **h**, The positioning of mismatched bases in each sgRNA targeted to EGFP site 1. **i**, Double mismatch tolerance of the two-component VECTRv2-Cas(sgGFP) vs. LentiCRISPRv2(sgGFP) harboring variant mismatched sgRNAs. A matched sgRNA for site 1 was used as a control. (**b, e, g, i**) Mean activities of three replicates are shown. Error bars, mean ± s.e.m.

Next, we used an EGFP disruption assay to determine whether the “two-component” lentiviral <u>ve</u>ctor for the <u>c</u>ombined <u>tr</u>ansduction of the <u>Cas</u>9 protein and U6-sgRNA expression cassette (henceforth referred to as VECTR-Cas), produced from the transfected cells, can deliver the Cas9 protein to the nucleus of mammalian cells, form a complex with nascent sgRNA and induce mutagenesis. We targeted a specific site in a single copy of the *EGFP* reporter gene incorporated into chromosome 17 of the genome of HEK293 cells (HEK293-EGFP; **Supplementary Fig. 2**). Transduction of the unconcentrated VECTR-Cas(sgGFP) resulted in a robust loss of EGFP in ~90% of the EGFP-expressing cells (**Fig. 1b; Supplementary Fig. 3a**).

Titration of the pVpr.Prot.Cas9 plasmid used to prepare VECTR-Cas(sgGFP) revealed that the greatest extent of reporter *EGFP* gene disruption (~98%) was achieved with 0.9 μg of the construct in a total amount of 4 μg of transfected DNA (**Supplementary Fig. 4**).

The level of disruption was almost as high as that obtained with the potent *cas9* gene-delivering lentiviral vector pLentiCRISPRv2(sgGFP) ^21^. The level of EGFP expression was unchanged after transduction of a control lentivector lacking the Vpr.Prot.Cas9 protein or containing an sgRNA that targeted an EMX1 locus (**Fig. 1b**; **Supplementary Fig. 3b**). No significant EGFP disruption was observed when the vector particles were prepared in the absence of the VSV.G envelope protein or when an inhibitor of reverse transcription, azidothymidine (AZT), was added to transduced cells (**Fig. 1b**; **Supplementary Fig. 3b**).

To confirm that the disruption of *EGFP* arose from genome modification and not from Cas9 binding or cellular toxicity, we measured the frequency of insertions/deletions (indels) at the target EGFP locus by means of a T7 endonuclease (T7E1) assay and Inference of CRISPR Edits (ICE) analysis (ice.synthego.com). Both assays revealed the presence of VECTR-Cas-induced double-strand breaks (DSBs) corrected by error-prone nonhomologous end joining (NHEJ). The frequencies of gene disruption differed (38% for T7E1 vs. 97% for ICE) (**Fig. 1c and d**; **Supplementary Fig. 5**), perhaps due to a high frequency of +1 nt insertions (37% of indels) that increased the likelihood of reannealing the mutant DNA strands, leading to insensitivity of the resulting homoduplexes to T7E1 (**Fig. 1c and d**; **Supplementary Fig. 5**) ^22^. In contrast to transfection, transduction did not cause any appreciable toxicity in HEK293-EGFP cells, as validated by an XTT assay that showed only a minimal loss of cell viability after transduction (**Supplementary Fig. 6**).

The use of an integration-deficient vector would prevent the possibility of integrating lentiviral DNA, thereby improving the safety of two-component nanoparticles. Efficient knockouts mediated by programmable nucleases require only a short burst of activity, so we reasoned that the expression of sgRNAs from unintegrated viral DNA might be sufficient for gene ablation. We therefore tested the ability of vector particles containing a catalytically inactive integrase protein (D64V, VECTRv2-Cas(sgGFP)) ^23^ to disrupt EGFP. There was no detectable difference in gene ablation activity compared with that of the integration-proficient VECTR-Cas(sgGFP), indicating that the integration-deficient vector is a viable modification of the system (**Fig. 1e**; **Supplementary Fig. 7**).

Transducing HEK293-EGFP cells with decreasing amounts of LVs revealed a positive correlation between the dose of lentiviral vector and the disruption of EGFP. The loss of knockout activity was more pronounced for the two-component VECTRv2-Cas(sgGFP) nanoparticles than for the gene-delivering pLentiCRISPRv2(sgGFP), consistent with the idea that direct protein delivery is more vulnerable to a reduction of the effective nuclease protein concentration than is the administration of a nuclease gene expression cassette (**Supplementary Fig. 8**).

We compared the kinetics of EGFP disruption between VECTRv2-Cas(sgGFP) and pLentiCRISPRv2(sgGFP) by following the expression of EGFP over an 80 h period after transduction. There was no difference in the time course of gene disruption. Both RGEN delivery methods required a lag period of over 24 h to disrupt EGFP, which was followed by a steady increase in disruption, plateauing at 56 h post transduction (**Fig. 1f**; Supplementary **Fig. 9a**). The loss of fluorescence remained stable over a period of 35 days after treatment (**Supplementary Fig. 9b**). In combination with the findings obtained with AZT, which indicated that reverse transcription is required for site-specific mutagenesis (**Fig. 1b**), these results support a model whereby a complex between Cas9 and sgRNA is formed in the nucleus of the recipient cells rather than during the assembly of viral particles (**Fig. 2**).

**Fig. 2:**
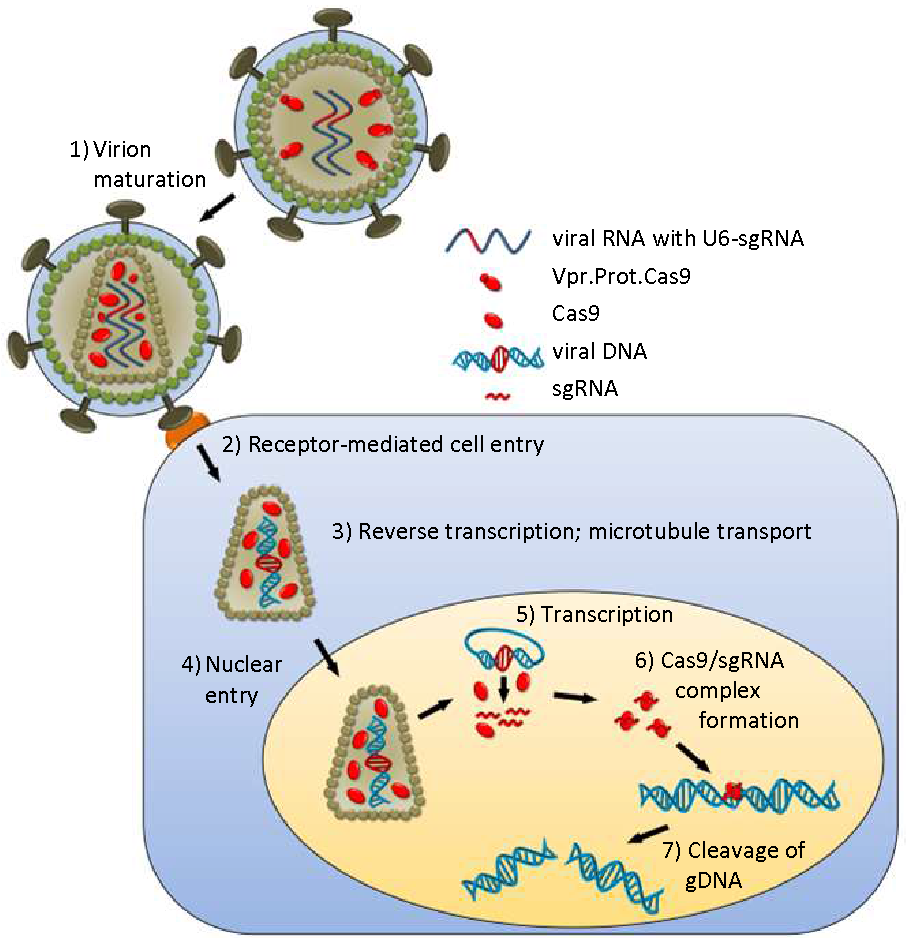
A schematic representation of lentivector-mediated delivery of the Cas9 protein and viral RNA containing U6-sgRNA. Cas9 is packaged into virions as a Vpr.Prot.Cas9 fusion polyprotein that is proteolytically cleaved during virion maturation (1). Following virus entry into a recipient cell (2), the viral genome is reverse transcribed to DNA (3) and translocated to the nucleus together with Cas9 (4), where the U6 promoter drives the expression of sgRNA (5). The nascent sgRNA associates with Cas9 (6) and directs the nuclease to the target site in the genomic DNA (gDNA) for cleavage (7).

To ensure that disruption of EGFP is not a peculiarity of the sgRNA used (sgGFP site No. 1), we tested the activity of a set of *EGFP*-targeting sgRNAs (sgGFP site Nos. 2–8). One of these sgRNAs (targeting site No. 2) induced indels with a frequency comparable to that mediated by the original sgRNA. The remaining six sgRNAs yielded mutation rates that ranged from 12% to 85% (**Fig. 1g**). The gene delivery vectors consistently exhibited more robust editing activity than the vectors that directly transferred the Cas9 protein, although the magnitude of the difference was not the same for all sites. The effect was particularly salient for the sgRNA that targeted site No. 8, which was the least edited site (**Fig. 1g**; **Supplementary Fig. 10**). It appears that some of the targeted loci are more sensitive to changes in Cas9/sgRNA concentrations in the cell and thus that the design of the sgRNAs is particularly important for systems that directly deliver RGENs to cells.

We next investigated whether the VECTRv2-Cas system is more sensitive than the lentivirus-mediated administration of nucleic acids to Watson-Crick mismatches at the sgRNA-DNA interface. We determined the EGFP disruption activity of both lentivirus-based delivery systems bearing variants of the original sgRNA (sgGFP site No. 1) with adjacent double mismatches at positions 1-19 (numbered in the 3’ to 5’ direction; **Fig. 1h**). Regardless of the route of RGEN administration, the EGFP disruption activity was robust and equivalent to that mediated by the matched sgRNA when the sgRNA contained mismatches at positions 17&18 and 18&19. The sgRNA containing mismatches at position 15&16 yielded less potent editing than the matched sgRNA, and the loss of activity was more dramatic when the sgRNA was transduced into cells together with the Cas9 protein. Of the remaining seven sgRNAs, only sgGFP(11&12) showed appreciable activity and only when delivered via the pLentiCRISPRv2 encoding the nuclease (**Fig. 1i**; **Supplementary Fig. 11**). These results are consistent with reports that mismatches at the 5’ end of sgRNAs are better tolerated than those at the 3’ end and establish that VECTR-Cas-mediated editing is more specific than mutagenesis induced by the transduction of the *cas9* gene into cells ^24,25^.

To test whether virions delivering Cas9 protein:template sgRNA can induce the cleavage of endogenous genes in human cells, we transduced HEK293-EGFP cells with VECTRv2 targeted to the EMX1, FANCF, HEKsite1 and HEKsite3 loci in the human genome. The T7E1 assay revealed that indels formed at all four genomic loci with efficiencies similar to those achieved with cas9 gene-delivering virions (13%–55%) (**Fig. 3a**; **Supplementary Fig. 12 and 13**). VECTRv2-Cas can also induce site-specific DSBs in more technically challenging cell types, including the Jurkat, SupT1, IM9, and THP-1 cell lines, as well as primary CD4+ T cells. Indels formed at frequencies ranging from 6% to 70% in all cell types tested (**Fig. 3b and c**; Supplementary **Fig. 14 and 15**). IM9, THP-1, and CD4+ T cells were less edited, but the two T cell-derived lines, Jurkat and SupT1, were as sensitive to the induction of mutations as HEK293-EGFP cells. We used amplicons generated with genomic DNA from SupT1 and CD4+ T cells for high-throughput sequencing to verify and quantify on-target mutations at the FANCF, HEKs1, and HEKs3 loci. For CD4+ cells, we found indels at frequencies of ~3%, 15%, and 13%, respectively, while transduction of SupT1 yielded cleavage efficiencies of 75%, 100% and 100%, respectively (**Fig. 3d and e**) (**Supplementary Fig. 16-18**).

**Fig. 3:**
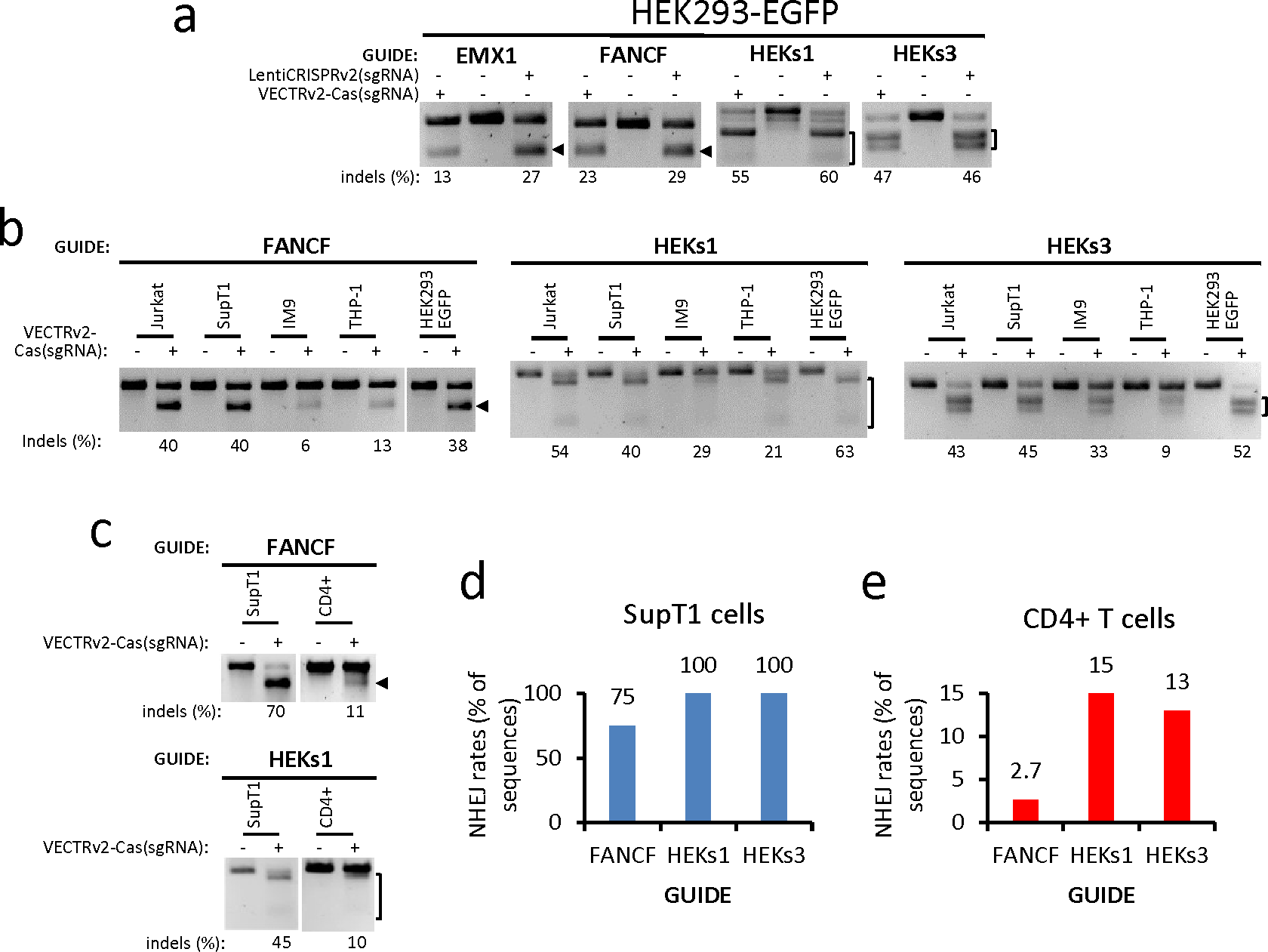
RNA-guided genome editing of the native loci in multiple cell types. **a**, Detection of indels by a T7E1 assay in the endogenous EMX1, FANCF, HEKs1, and HEKs3 loci in HEK293-EGFP cells transduced with VECTRv2-Cas(sgRNA) or LentiCRISPRv2(sgRNA). **b**, VECTRv2-Cas(sgRNA)-mediated mutation of the FANCF, HEKs1 and HEKs3 loci in lines of human T lymphocytes (Jurkat, SupT1), B lymphocytes (IM9) and monocytes (THP-1) measured with the T7E1 assay. Mutation rates obtained from parallel transductions of HEK293-EGFP cells are also shown. **c**, NHEJ rates (measured by the T7E1 assay) in primary CD4+ T cells transduced with VECTRv2-Cas(sgRNA). Parallel transduction of SupT1 cells served as a positive control. Please note that the HEKs3 site could not be analyzed by the T7E1 assay due to a single-nucleotide polymorphism (SNP) near the cut site (**Supplementary Fig. 17**). **d-e,** NHEJ frequencies quantified by CRISPR Genome Analyzer using next-generation sequencing data of amplicons from (**c**) as an input ^26^. **d**, SupT1 T cell line; **e**, primary CD4+ T cells.

Here, we describe a novel system for the codelivery of the Cas9 protein and a template for sgRNA within lentivirus-based “nanoparticles”. The idea is built upon findings that lentiviruses can simultaneously deliver foreign proteins and an episomal viral DNA generated by reverse transcription from the vector RNA genome ^9,10^. We have extended this approach to show that episomal DNA can serve as a template for the transcription of sgRNA, which forms a complex with the co-delivered Cas9 protein and targets the nuclease to a specific site in the genome. This strategy leads to robust editing activity that is comparable or even superior to that reported for the direct delivery of Cas9 protein/sgRNA complexes to cells ^1,2^. In contrast to chemical transfection or electroporation, virus-mediated delivery is receptor mediated, so the use of pseudotypes bearing natural or engineered envelope proteins would allow selective transfer to essentially any target cell population ^6^. This approach may even extend the repertoire of cell types that can be edited to clinically relevant nondividing cells (e.g. neurons, hepatocytes, quiescent lymphocytes and hematopoietic stem cells) that are permissive for the lentivector-mediated transduction of cargos to the nucleus ^5^. Additionally, because the removal of cas9 transgene from the lentiviral transfer vector makes a space for additional function-conferring elements, we imagine that the approach can be developed to a multiple-component system for simultaneous knock-out and delivery of genes. For example, simultaneous disruption of the T cells receptor (TCR) and delivery of the chimeric artificial receptor (CAR) may simplify the development and manufacturing of the advanced medicinal products such as allogenic CAR-T cells.

There are several possible explanations for the high efficiency of the two-component VECTRv2-Cas nanoparticles. First, we fused Cas9 to Vpr rather than to the Gag polyprotein, as we anticipated that the incorporation of the bulky nuclease into Gag would destabilize the virions and lower the efficiency of transduction. Second, we incorporated the RRE into the transcripts encoding Gag and pVpr.Prot.Cas9 proteins, providing simultaneous control of transport and spatial distribution to both mRNAs and resulting in the accumulation of Vpr.Prot.Cas9 in viral particles. ^12,13^. This probably results from the colocalization of nascent Gag and Vpr.Prot.Cas9 proteins after translation, facilitating the interaction between p6 (in Gag) and Vpr (in Vpr.Prot.Cas9) and stimulating the incorporation of the fusion protein into virions (**Supplementary Fig. 1b**). Alternatively, analogous post-transcriptional regulation of gene expression might have allowed the interaction with a cellular factor responsible for the trafficking of transcripts and/or proteins to the virus assembly site. Third, the Cas9 protein remains embedded in a lentiviral core after receptor-mediated cell entry (where it is protected from degradation) and exploits viral intracellular trafficking routes to travel to the nucleus. Finally, the coordinated intranuclear delivery of Cas9 and viral DNA within a preintegration complex particle probably places the nuclease in close proximity to nascent sgRNA molecules synthesized from viral DNA, facilitating the formation of Cas9/sgRNA RNP complexes (**Fig. 2**).

In summary, we describe novel multicomponent lentiviral nanoparticles that ferry the Cas9 protein:sgRNA template to the nuclei of transduced cells for the transient exposure of the genome to the nuclease that results in the specific disruption of targeted genes. This system may represent a versatile platform for the efficient, safe, nontoxic and cell type-selective delivery of genome modification enzymes to cells.

## Supporting information

Supplementary_figures_methods_references

## Data Availability Statement

Raw MiSeq sequencing files are publicly available at the European Nucleotide Archive (ENA; www.ebi.ac.uk/ena) under accession number: PRJEB32556.

## Acknowledgements

This work was supported by the Austrian Science Fund (FWF) to SI [P24577]. The funders had no role in study design, data collection and analysis, decision to publish, or preparation of the manuscript.

## Author contributions

I.I., Conception and design, Acquisition of data, Analysis and interpretation of data, Revising the article; S.I., Conception and design, Acquisition of data, Analysis and interpretation of data, Drafting and revising the article.

## Competing interests

S.I. and I.I. have filed a patent related to this work.

