## Supplementary_figures_methods_references for "A novel toolkit for the efficient delivery of Cas9/sgRNA complexes to chromosomes in cells"

#### SUPPLEMENTARY INFORMATION


| | | |
|---|---|---|
| **Supplementary figure 1** | Enhanced packaging of the Vpr.Prot.Cas9 fusion protein when the expression cassette encodes a transcript that comprises the Rev responsive element (RRE) | 2-3 |
| **Supplementary figure 2** | Production of reporter HEK293-EGFP cells | 4 |
| **Supplementary figure 3** | Transduction of bi-component lentivector containing Cas9 protein and U6-sgRNA cassette disrupts EGFP gene. | 5 |
| **Supplementary figure 4** | Titration of the pVpr.Prot.Cas9 plasmid for optimal gene disruption activity. | 6 |
| **Supplementary figure 5** | Determination of EGFP gene editing efficiency by Inference of CRISPR Edits | 7 |
| **Supplementary figure 6** | Transduction with two-component lentivector has minimal effect on cell viability. | 8 |
| **Supplementary figure 7** | Integration deficient vector mediates EGFP disruption as efficiently as the vector containing wild-type (WT) integrase (IN). | 9 |
| **Supplementary figure 8** | EGFP disruption assay performed with decreasing doses of vectors. | 10 |
| **Supplementary figure 9** | Time course of EGFP knockout after transduction with the two-component Cas9 protein-containing VECTRv2-Cas or *cas9* gene-carrying lentivector (LentiCRISPRv2(sgGFP)). | 11 |
| **Supplementary figure 10** | EGFP disruption activity of Cas9 protein- and *cas9* gene-carrying lentivectors. | 12 |
| **Supplementary figure 11** | EGFP disruption activity of Cas9 protein- and *cas9* gene-carrying lentivectors containing either matched sgRNA (site 1) or sgRNA bearing double adjacent mismatches at the indicated positions. | 13 |
| **Supplementary figure 12** | Determination of editing efficiency in the HEKs1 locus by Inference of CRISPR Edits after transduction with VECTRv2-Cas(sgHEKs1) | 14 |
| **Supplementary figure 13** | Determination of editing efficiency in the HEKs1 locus by Inference of CRISPR Edits after transduction with LentiCRISPRv2(sgHEKs1). | 15 |
| **Supplementary figure 14** | Determination of editing efficiency in the FANCF locus by Inference of CRISPR Edits after transduction with VECTRv2-Cas(sgFANCF) | 16 |
| **Supplementary figure 15** | Determination of editing efficiency in the FANCF locus by Inference of CRISPR Edits after transduction with LentiCRISPRv2(sgFANCF). | 17 |
| **Supplementary figure 16** | Frequency of VECTRv2-Cas(sgFANCF)-mediated indel formation by non-homologous end joining (NHEJ) in lymphocytes. | 18 |
| **Supplementary figure 17** | Frequency of VECTRv2-Cas(sgHEKs1)-mediated indel formation by non-homologous end joining (NHEJ) in lymphocytes. | 19 |
| **Supplementary figure 18** | Frequency of VECTRv2-Cas(sgHEKs3)-mediated indel formation by non-homologous end joining (NHEJ) in lymphocytes. | 20 |
| **Material and methods** | | 21-23 |
| **List of oligos** | | 24-25 |
| **Supporting references** | | 26-27 |

#### SUPPLEMENTARY FIGURE 1

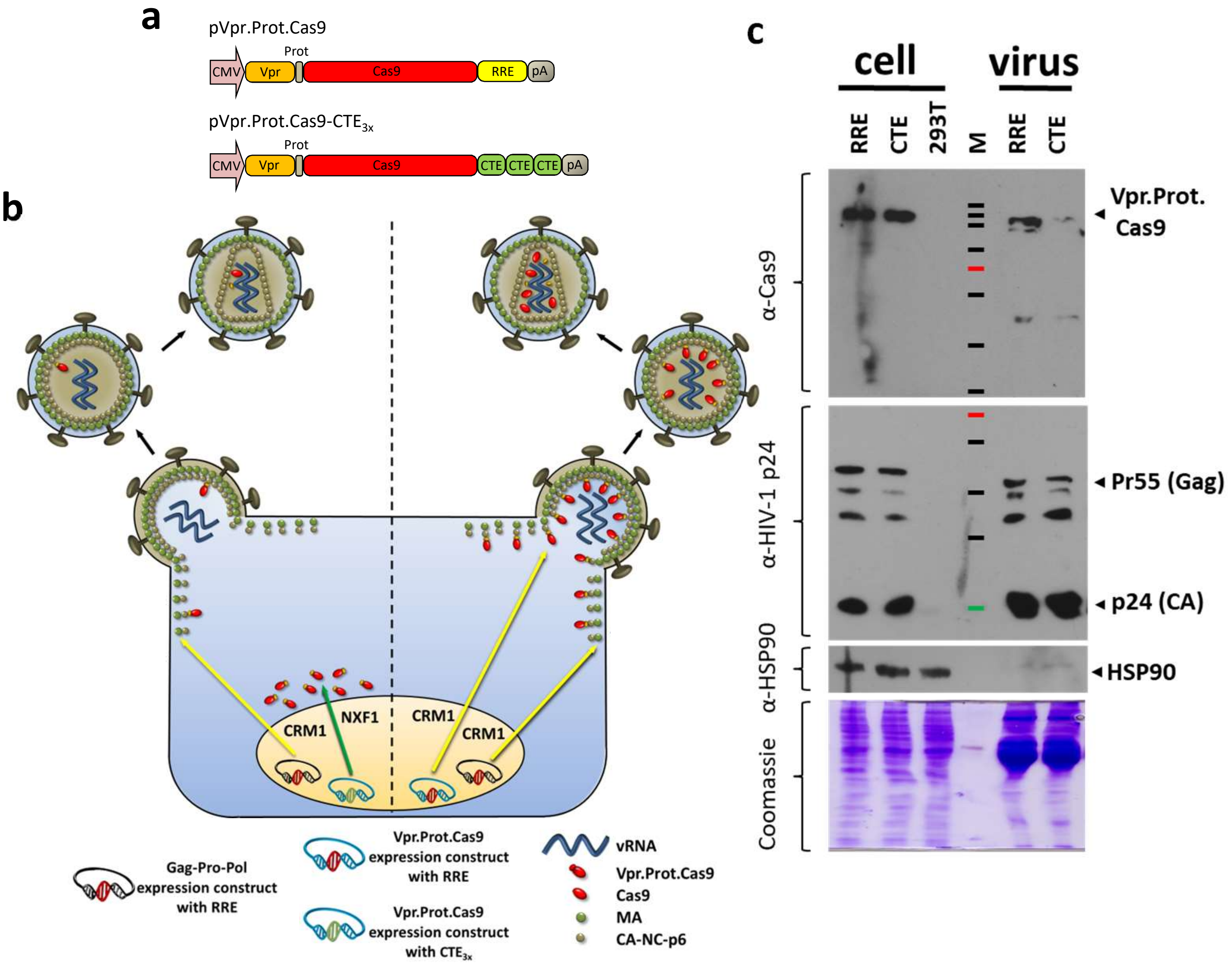


**Supplementary figure 1 | Enhanced packaging of the Vpr.Prot.Cas9 fusion protein when the expression cassette encodes a transcript containing the Rev responsive element (RRE).**

(**a**) The expression cassettes encoding Vpr.Prot.Cas9 fusion protein (**b**) Retrovirus assembly and budding is a highly concerted process. It is mediated by numerous, largely undefined spatially and temporally regulated interactions between viral proteins and cellular factors. Previous reports showed that the regulation of the HIV-1 Gag assembly begins as soon as nuclear export factors are deposited onto the transcripts encoding the structural components of HIV-1. These studies are consistent with the model that selection of the RNA export pathway modulates the cytosolic fate and function of viral transcripts and/or structural polyproteins (Swanson, C. M., et al. (2004); Jin, J., et al. (2007); Sherer, N. M., et al. (2009); Pocock, G. M., et al., 2016). We hypothesized that the RNA export pathway selection will also analogously affect the cytosolic fate of the non-viral transcripts and/or heterologous proteins. Further, we postulated that nuclear export of both viral and non-viral transcripts via the same pathway will facilitate the cytoplasmic co-localization of the transcripts and their translation products. A close proximity of Gag and Vpr.Prot.Cas9 promotes the interaction between the two polyproteins that is required for packaging of the fusion protein to virions. To test this hypothesis, we constructed two Vpr.Prot.Cas9 expression constructs carrying various RNA export functions, Vpr.Prot.Cas9 (contains Rev responsive element, RRE) and Vpr.Prot.Cas9-$CTE_{3x}$ (contains constitutive transport element from MPMV; three copies of CTE were used to produce approximately the same protein levels in the cytoplasm (Wodrich, H., et al., 2000)). The RRE facilitates mRNA export via the CRM1 pathway, whereas the CTE drives mRNA nuclear exit via the NXF1 pathway (Braun, I. C., et al., 1999; Fornerod, M., et al., 1997; Fukuda, M., et al., 1997; Stade, K., et al., 1997). (**b**) The

**Continuation of Supplementary figure 1 |**
constructs were co-transfected into HEK293T cells together with psPAX2 (encodes structural and enzymatic components of virions and the transcript contains RRE), and with the other plasmids to generate viruses as described for Fig.1a. The presence of viral and heterologous proteins in the cell lysates (cell, 10 µg of total protein) and virions harvested from cell culture supernatant 48 hours after transfection (virus, 100x concentrated by ultracentrifugation [20 000 g, 4°C, 2h]) was determined by immunobloting with the antibodies as described below. The immunoblot obtained with an antibody specific for Cas9 protein (α-Cas9 [mouse, 1:200, 7A9-3A3; Santa Cruse]) showed the same levels of Vpr.Prot.Cas9 protein in the cell lysates of cells transfected with either of the two Vpr.Prot.Cas9 expression constructs (cell). In sharp contrast, the fusion protein could be detected in the virus pellets only when the transcript encoding Vpr.Prot.Cas9 contained the RRE (virus). This result was not due to a differential loading or blotting as the same levels of HIV-1 proteins (p24 and Pr55) were detected in both virus preparations following re-probing of the membrane with an antibody raised against HIV-1 capsid protein (α-HIV-1 p24, rabbit, 1:2000, a gift from Dr. Sakalian). Equivalent expression of the viral proteins (p24 and Pr55) was also found in the cell lysate preparations containing the same levels of the loading control cellular protein HSP90 (α-HSP90 [rabbit, 1:1000, Cell Signaling Technology]).

Collectively, these results support the view that the RNA nuclear export pathway selection governs transport dynamics and/or subcellular localization of viral and heterologous mRNAs. Matching spatial localization and/or temporal distribution of the *gag* mRNA and *vpr.prot.cas9* transcripts, and in turn the Gag and Vpr.Prot.Cas9 proteins, promotes the interaction between the two nascent polypeptides and results in increased levels of the fusion protein in virus particles (**a**).

The α-rabbit-IgG-HRP (1:10000, DAKO, P0399) and α-mouse-IgG-HRP (1:2000, DAKO, P0260) secondary antibodies were used. M: Protein Standard (NEB#P7719). Proteins in the SDS-PAGE gel were stained with Coomassie dye after blotting. One representative example from two biological replicates performed in two different weeks is shown.

#### SUPPLEMENTARY FIGURE 2

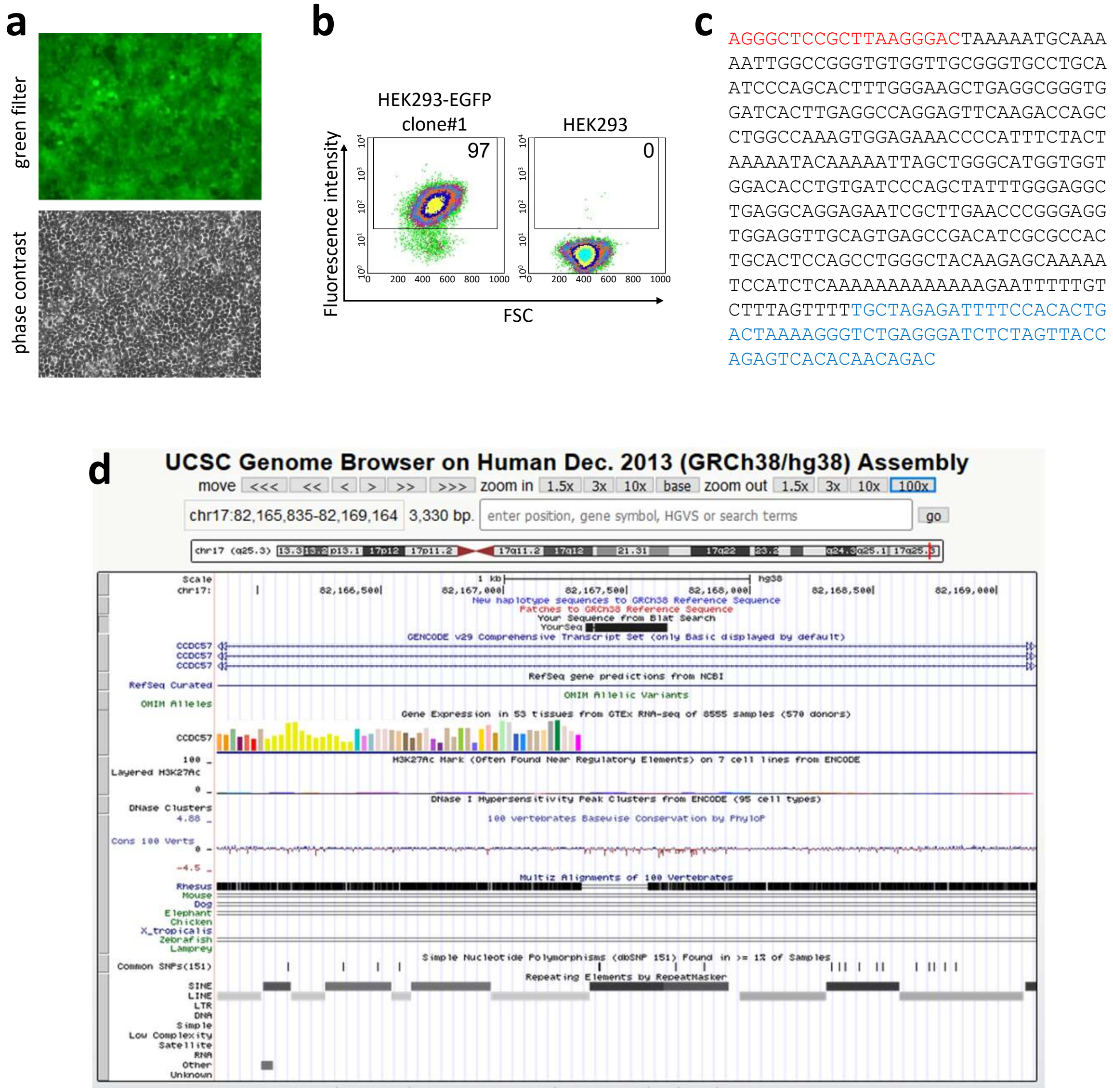


**Supplementary figure 2 | Production of reporter HEK293-EGFP cells**

HEK293-EGFP cells were prepared by clonal selection of HEK293 cells transduced with a lentiviral vector prepared by co-transfection of three plasmids: i) a transfer vector containing a CMV-EGFP expression cassette and a hygromycin resistance gene (pLenti CMV GFP Hygro, addgene #17446); ii) a packaging construct psPAX2 (addgene #12260); iii) a plasmid encoding the VSV-G envelope (pHCMV-G). A low multiplicity of infection (MOI = 0.05) was used to ensure a single integration event per cell. Following transduction, the cells were subjected to cloning by limiting dilution. Monoclonal cell populations exhibiting a minimal mean fluorescence intensity (MFI ~ 100), as determined by UV microscopy (**a**) and flow cytometry (FACS Calibur) (**b**), were passaged for approximately 30 weeks. The clones showing a stabile and homogenous EGFP expression were then used for the determination of integration sites of the lentivector by a ligation-mediated PCR (LM-PCR) (Wu et al., 2003). The clone #1, containing the lentivector DNA integrated at the chromosome 17, as determined by BLAT search algorithm, was used for further experimentation (**c and d**). (**c**) Integration site as determined by Sanger sequencing of the LM-PCR product (linker sequence is shown in red, human sequence in black and vector sequence in blue color).

#### SUPPLEMENTARY FIGURE 3

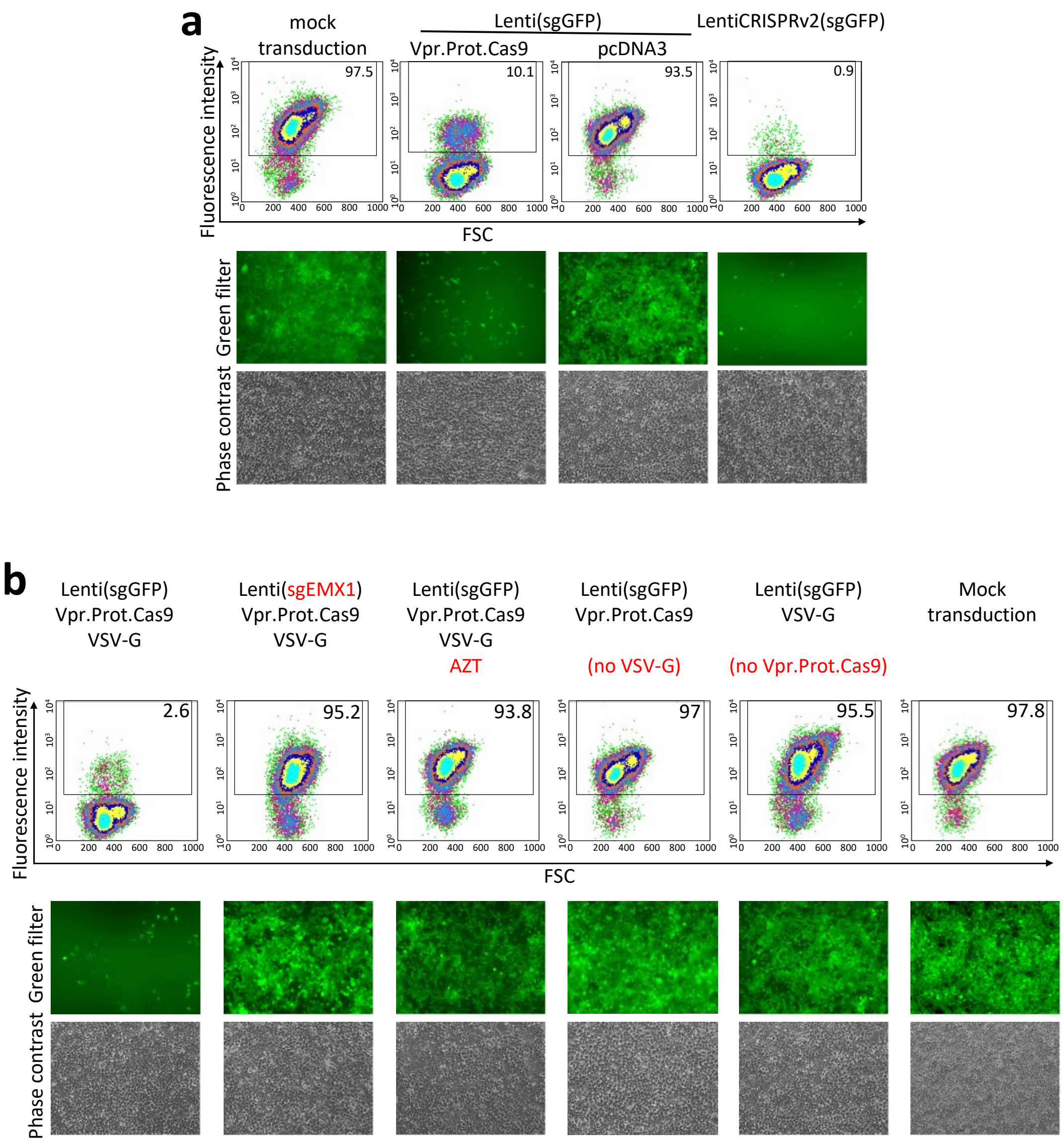


**Supplementary figure 3 | Transduction of two-component lentivector containing Cas9 protein and U6-sgRNA cassette disrupts EGFP gene. Additional data to Fig. 1b.** (**a**) Flow cytometry analysis (dot plots in the upper panel) and UV microscopy (lower panels) performed three days post transduction revealed an efficient disruption of the *EGFP* gene when pVpr.Prot.Cas9 was present during vector preparations. (**b**) Flow cytometry (upper panels) and UV microscopy (lower panels) demonstrated that the EGFP disruption is abolished when (i) sgRNA specific to the EMX1 locus was cloned into pLenti(sgRNA) in place of the sgRNA targeting EGFP; (ii) an inhibitor of reverse transcription (azidothymidine, AZT) was added to the culture medium; (iii) the VSV-G expression construct pHCMV.G or (iv) the pVpr.Prot.Cas9 plasmid was omitted during vector preparation. (**a**,**b**) Representative images from three replicates are shown. The percentage of EGFP positive cells is shown in the upper right corner of the flow cytometry dot blots.

#### SUPPLEMENTARY FIGURE 4

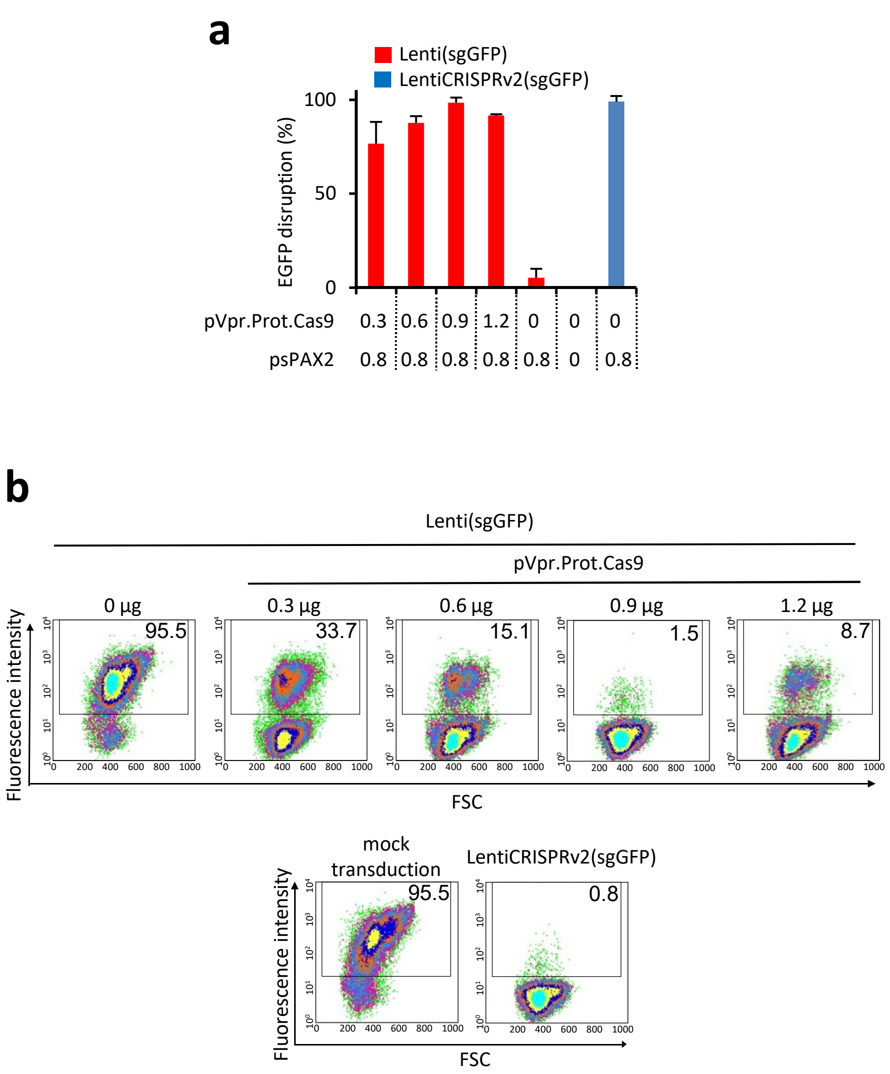


**Supplementary figure 4| Titration of the pVpr.Prot.Cas9 plasmid for optimal gene disruption activity.** (**a**) HEK293T cells were transfected with a total amount of 4 µg of plasmid DNA. The amount of the Cas9 expression construct varied from 0 µg to 1.2 µg (adjusted to 1.2 µg by "empty" pcDNA3). As a positive control LentiCRISPRv2(sgGFP) prepared in the absence of pVpr.Prot.Cas9 was used. Error bars reflect SD from three replicates. (**b**) representative examples of flow cytometry results (dot blots) used to calculate mean EGFP disruption activity for (**a**). The percentage of EGFP positive cells is shown in the upper right corner.

#### SUPPLEMENTARY FIGURE 5

**a**

HEK293-EGFP cells

| Label | ICE (indels %) | KO-Score * | R2 ** |
|---|---|---|---|
| VECTR-Cas(sgGFP) | 97 | 73 | 0.98 |

* Proportion of indels that indicate a frameshift or are 21+bp in length

** How well the proposed indel distribution fits the Sanger sequence data of the edited sample

**b**

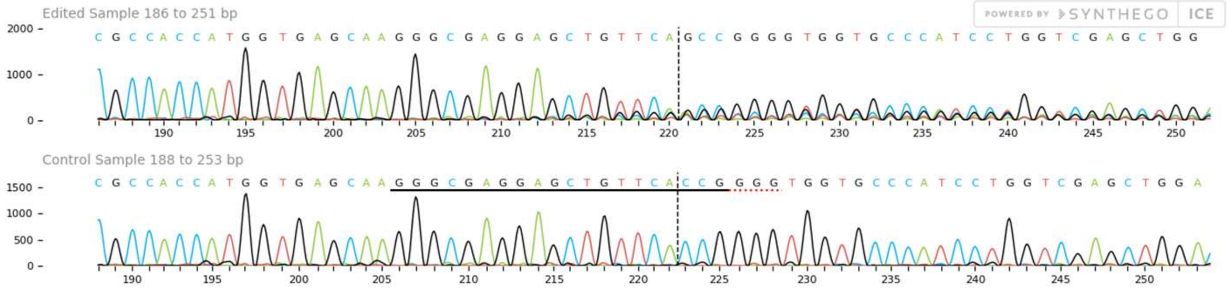


**c**

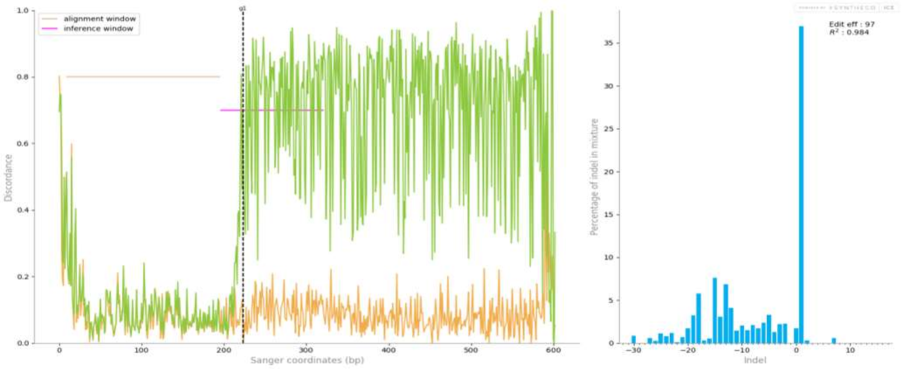


**d**

| contribution | indel | sequence (left of cut) | sequence (right of cut) |
|---|---|---|---|
| | wt | GTGAGCAAGGGCGAGGAGCTGTTCA | CCGGGTGGTGCCCATCCTGG (PAM: GGG) |
| 36.96% | +1 | GTGAGCAAGGGCGAGGAGCTGTTCA | nCCGGGGTGGTGCCCATCCTG |
| 6.25% | -15 | GTGAGCAAGGGCGAGGAGCTGTTC- | --------------ATCCTGG |
| 3.93% | -13 | GTGAGCAAGGGCGAGGAGC------ | -------GGTGCCCATCCTGG |
| 3.34% | -18 | GTGAGCAAGGGCGAGGAG------- | -----------CCCATCCTGG |
| 3.28% | -5 | GTGAGCAAGGGCGAGGAGCTG---- | -CGGGGTGGTGCCCATCCTGG |
| 2.43% | -14 | GTGAGCAAGGGCGAGGAGCTG---- | ----------GCCCATCCTGG |
| 2.37% | -12 | GTGAGCAAGGGCGAGGA-------- | ----GGTGGTGCCCATCCTGG |
| 2.28% | -2 | GTGAGCAAGGGCGAGGAGCTGTT-- | CCGGGGTGGTGCCCATCCTGG |
| 2.23% | -3 | GTGAGCAAGGGCGAGGAGCTGTT-- | -CGGGGTGGTGCCCATCCTGG |
| 1.91% | -19 | GTGAGCAAGGGCGAGGA-------- | -----------CCCATCCTGG |
| 1.76% | -12 | GTGAGCAAGGGCGA----------- | -CGGGGTGGTGCCCATCCTGG |
| 1.73% | 0 | GTGAGCAAGGGCGAGGAGCTGTTCA | CCGGGGTGGTGCCCATCCTGG |
| 1.72% | -7 | GTGAGCAAGGGCGAGGAGCT----- | --GGGGTGGTGCCCATCCTGG |
| 1.71% | -6 | GTGAGCAAGGGCGAGGAGCTG---- | --GGGGTGGTGCCCATCCTGG |
| 1.59% | -13 | GTGAGCAAGGGCGAGGA-------- | -----GTGGTGCCCATCCTGG |
| 1.54% | -8 | GTGAGCAAGGGCGAGGAG------- | -CGGGGTGGTGCCCATCCTGG |
| 1.53% | -10 | GTGAGCAAGGGCGAGGAGCT----- | -----GTGGTGCCCATCCTGG |
| 1.52% | -18 | GTGAGCAAGGGCGAGGAGCTGT--- | ---------------TCCTGG |
| 1.5% | -9 | GTGAGCAAGGGCGAGGA-------- | -CGGGGTGGTGCCCATCCTGG |
| 1.42% | -11 | GTGAGCAAGGGCGAGGAGCTG---- | -------GGTGCCCATCCTGG |
| 1.35% | -13 | GTGAGCAAGGGCGAGGAGCT----- | --------GTGCCCATCCTGG |
| 1.28% | -4 | GTGAGCAAGGGCGAGGAGCTG---- | CCGGGGTGGTGCCCATCCTGG |
| 1.11% | -15 | GTGAGCAAGGGCGA----------- | ----GGTGGTGCCCATCCTGG |
| 1.11% | -25 | GTGAGCAAGGGCGA----------- | --------------ATCCTGG |
| 0.96% | -23 | GTGAGCAAGG--------------- | --------GTGCCCATCCTGG |

**Supplementary figure 5 | Determination of *EGFP* gene editing efficiency by Inference of CRISPR Edits (ICE from SYNTHEGO). Additional data to Fig. 1d.** An amplicon obtained from HEK293-EGFP cells transduced with bi-component VECTR-Cas(sgGFP) was analyzed by ICE. PCR product from mock-transduced cells was used as a control. Additional data to Fig. 1D (**a**) Summary of editing results. (**b**) Sanger sequencing chromatograms of the edited (upper panel) and control (lower panel) samples. The horizontal black underlined region represents the guide region. The PAM is underlined in red. The cut site is is shown by the vertical dotted line. (**c**) The left panel shows the level of disagreement between the control and edited samples around the cut site. Distribution of indel sizes and their frequencies is depicted in the right panel. (**d**) The contribution tab shows the sequences present in the pool of edited sequences. Their relative abundance is shown on the left. The cut site is represented by the red vertical dotted line.

#### SUPPLEMENTARY FIGURE 6

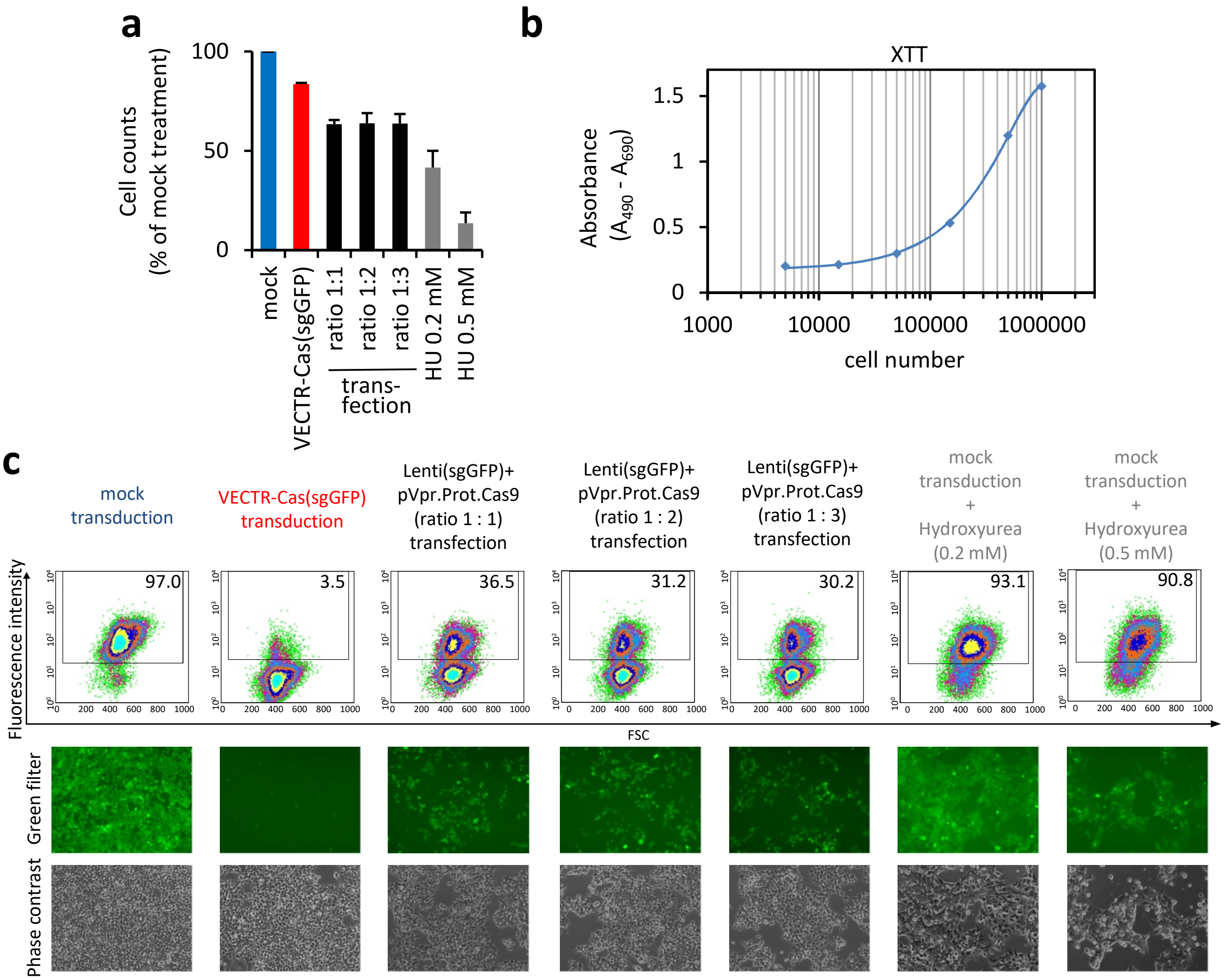


**Supplementary figure 6 | Transduction with two-component lentivector has a minimal effect on cell viability.** (**a**) Cell viability was measured by the Cell Proliferation (XTT) assay (Roche) three days post transduction with VECTR-Cas(sgGFP). HEK293-EGFP cells transduced in 12 well plates were incubated with yellow XTT solution (final concentration 0.3 mg/ml; 2 h) added directly to the cell culture media. Next, formation of orange formazan dye in cell culture medium by viable cells was quantified by an ELISA reader. As controls mock transduced cells, cells co-transfected with Lenti(sgGFP) + pVpr.Prot.Cas9 plasmids at a ratio of 1 : 1, 1 : 2 or 1 : 3 (Fugene HD; Promega); or cells treated with hydroxyurea (HU; 0.2 or 0.5 mM) were used. Mean viabilities of two biological replicates are shown. Error bars, mean ± SD. (**b**) Quantification of formazan dye formed from XTT tetrazolium salt added to different amounts of cells seeded in 12 well plates. Mean absorbance values ± SD from two replicates are shown. (**c**) To verify efficient disruption of EGFP gene following transduction, the HEK293-EGFP cells were also monitored by flow cytometry (upper panels) and UV microscopy (lower panels). Representative images from one of the cell proliferation experiments used to calculate means in (**a**) is shown. The percentage of EGFP positive cells is shown in the upper right corner of the flow cytometry dot blots.

#### SUPPLEMENTARY FIGURE 7

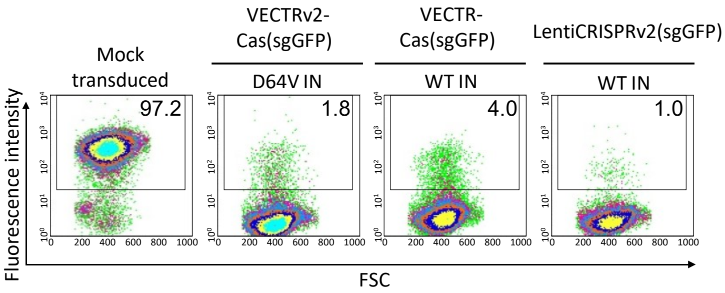


**Supplementary figure 7 | Integration deficient vector mediates EGFP disruption as efficiently as the vector containing wild-type (WT) integrase (IN). Additional data for Fig. 1e.** The expression of sgRNA from episomal non-integrated viral DNA forms is sufficient for a high level gene disruption. For this experiment viral vectors were prepared with either psPAX2 (WT IN; VECTR-Cas(sgGFP)) or psPAX2D64V (D64V IN; VECTRv2-Cas(sgGFP)) packaging construct. The psPAX2D64V encodes an integrase with the D64V mutation in the active center that abolishes integration activity of the enzyme. Representative examples of flow cytometry dot blots from three replicates are shown. LentiCRISPRv2(sgGFP) containing WT IN (encoded from psPAX2 plasmid) served as positive control. The percentage of EGFP positive cells is shown in the upper right corner.

#### SUPPLEMENTARY FIGURE 8

**a**

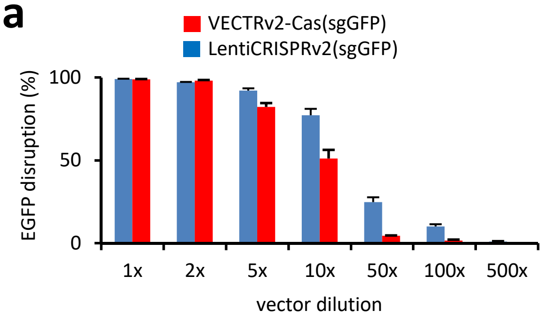


**b**

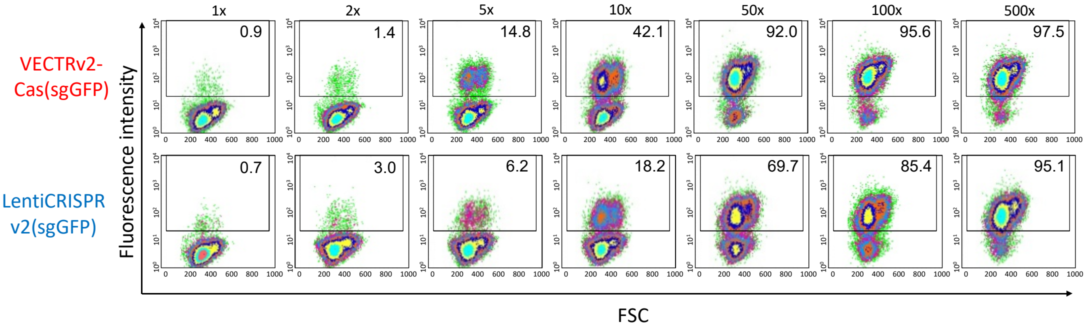


**Supplementary figure 8 | EGFP disruption assay performed with decreasing doses of vectors.** (**a-b**) Vector stocks were diluted in cell culture medium and subsequently used to transduce HEK293-EGFP cells. Flow cytometry, used to determine the EGFP disruption efficiency, was performed three days post transduction. The experiment was repeated twice and the dot blot figures in (**b**) represent representative examples from one of the experiments. The percentage of EGFP positive cells is shown in the upper right corner. (**a**) means ± SD.

#### SUPPLEMENTARY FIGURE 9

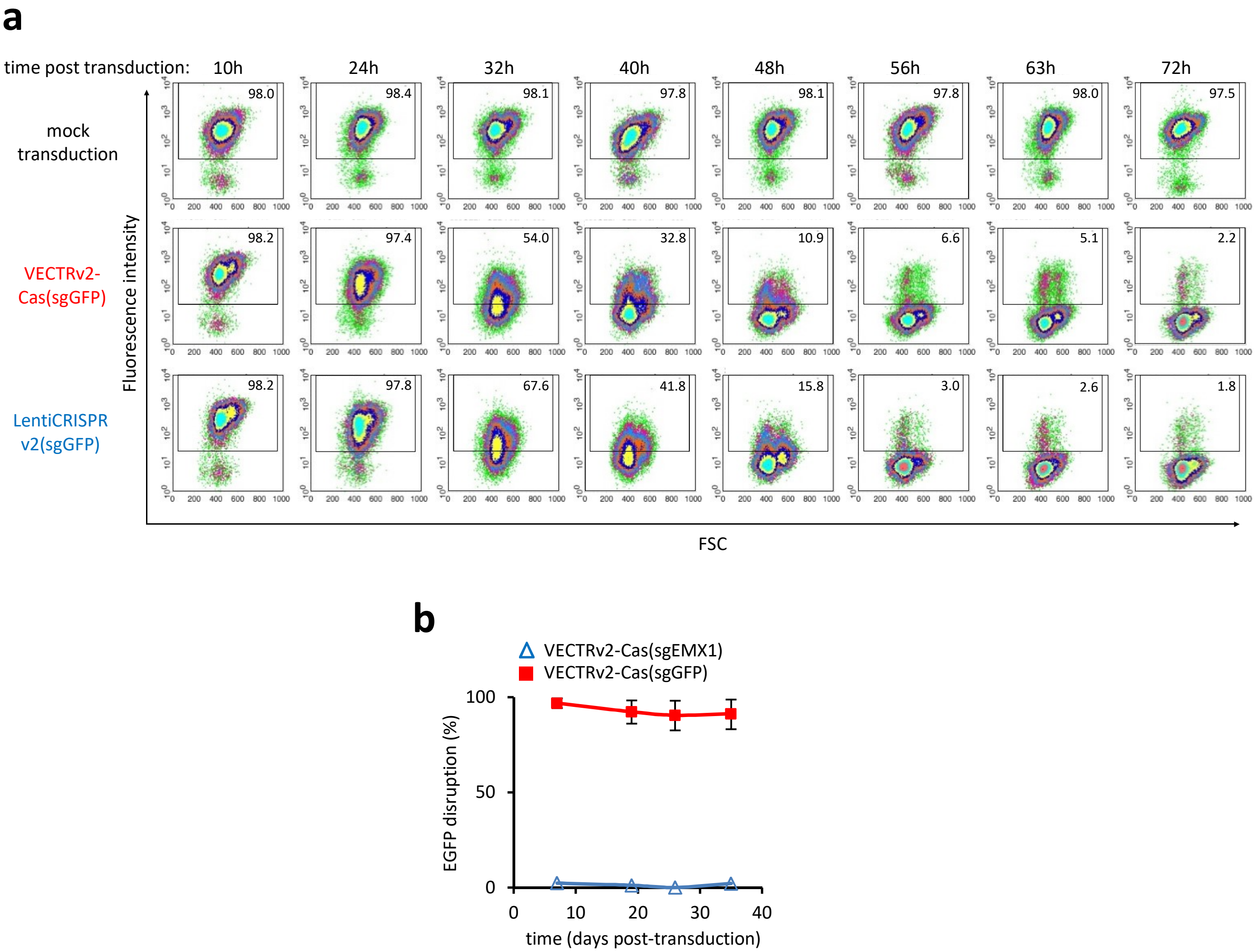


**Supplementary figure 9 | Time course of EGFP knockout after transduction with the two-component Cas9 protein-containing VECTRv2-Cas or *cas9* gene-carrying lentivector (LentiCRISPRv2(sgGFP)). Additional data for Fig. 1f.** (**a**) Flow cytometry was used to determine the percentage of EGFP-expressing cells at various time points after transduction that was then used for the calculation of disruption activity for the Fig. 1f. Dot blots of one of two replicates are shown. The percentage of EGFP positive cells is shown in the upper right corner. (**b**) Long term EGFP disruption experiment. EGFP expression in HEK293EGFP cells transduced with EGFP- or EMX1-targeting bi-component VECTRv2 was measured over a period of 35 days post transduction. The EGFP disruption values reflect means ± SD from four replicates.

#### SUPPLEMENTARY FIGURE 10

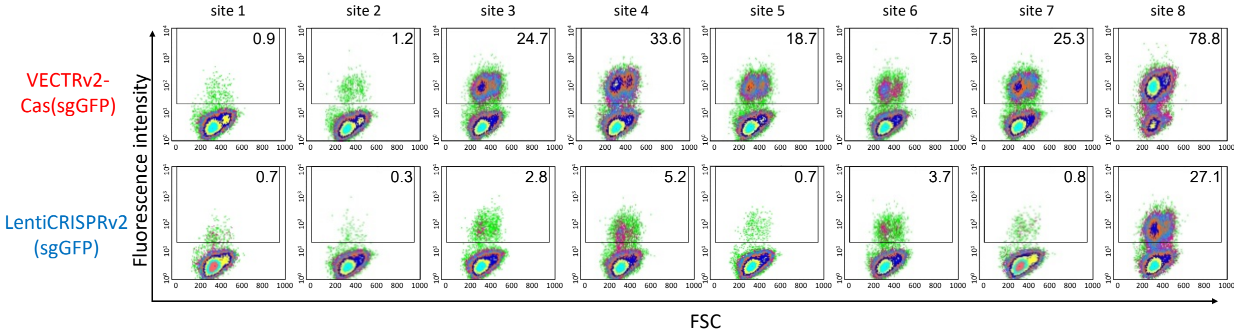


**Supplementary figure 10 | EGFP disruption activity of Cas9 protein- and *cas9* gene-carrying lentivectors targeting different loci in the *egfp* gene. Additional data to Fig. 1g.** The lentivector genomes contained U6 promoter driving the expression of sgRNAs directed against eight different loci within *egfp* gene (site 1- site 8). The decline of EGFP expression was measured by flow cytometry at day three post-transduction. Representative dot blots from one of three replicates used to calculate the disruption activities in Fig 1g are shown. The percentage of EGFP positive cells is shown in the upper right corner.

#### SUPPLEMENTARY FIGURE 11

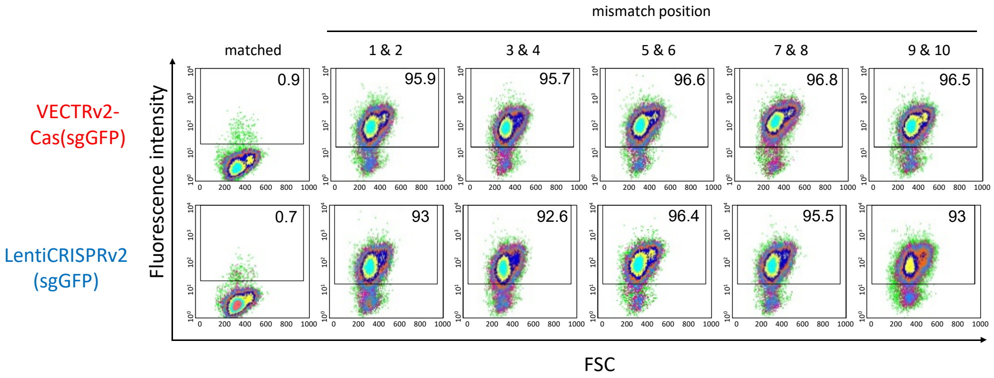


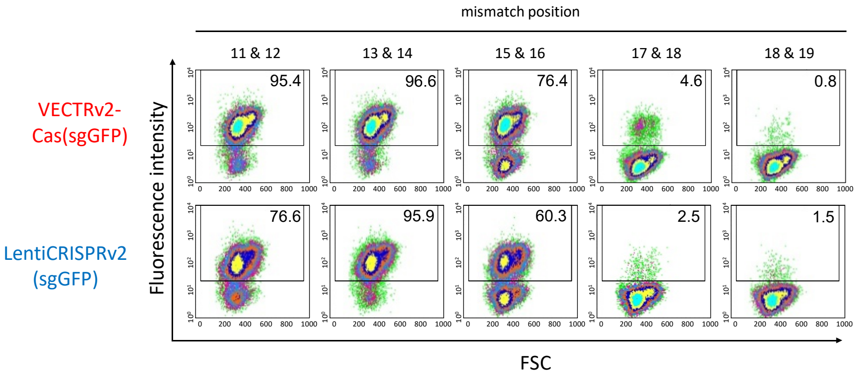


**Supplementary figure 11 | EGFP disruption activity of Cas9 protein- and *cas9* gene-carrying lentivectors containing either matched sgRNA (site 1) or sgRNA bearing double adjacent mismatches at the indicated positions. Additional data for Fig. 1h and i.** The flow cytometry dot blots reflect representative examples of three replicates used to calculate mean ± SD EGFP disruption activity shown in Fig 1i. The percentage of EGFP positive cells is shown in the upper right corner of the dot blots.

#### SUPPLEMENTARY FIGURE 12

**a**

HEK293-EGFP cells

| Label | ICE (% indels) | KO-Score * | $R^2$ ** |
|---|---|---|---|
| VECTRv2-Cas(sgHEKs1) | 99 | 88 | 0.99 |

* Proportion of indels that indicate a frameshift or are 21+bp in length
** How well the proposed indel distribution fits the Sanger sequence data of the edited sample

**b**

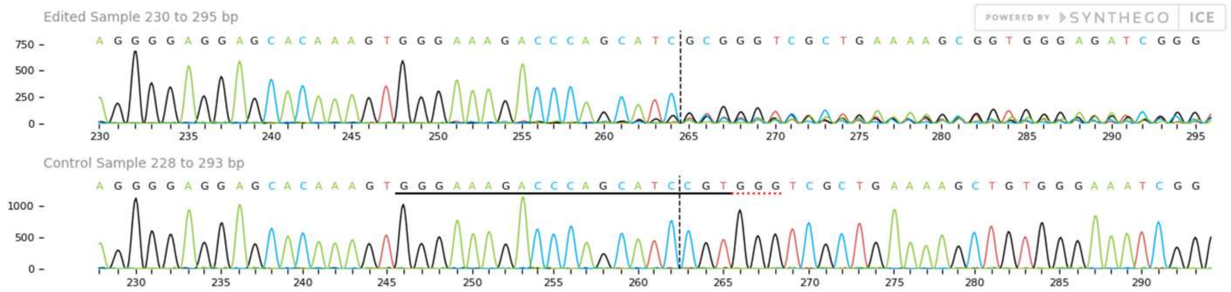


**c**

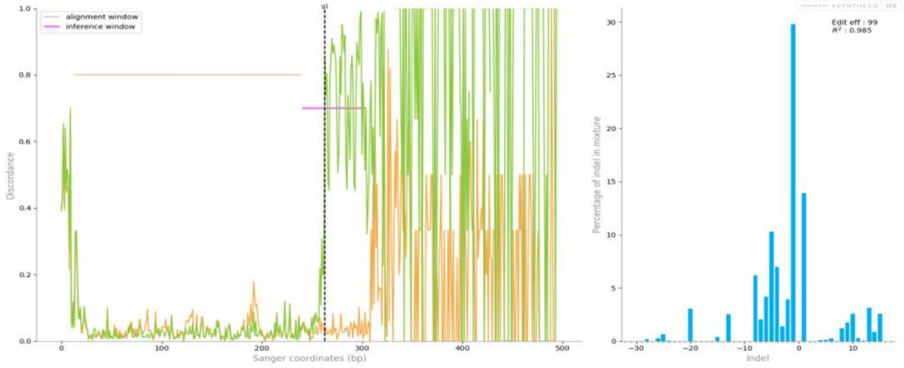


**d**

PAM

wt C A C A A A G T G G G A A A G A C C C A G C A T C | C G T G G G T C G C T G A A A A G C T G T G G G A A A T C G G G

| contribution | indel | sequence |
|---|---|---|
| 29.8% | -1 | C A C A A A G T G G G A A A G A C C C A G C A T - \| C G T G G G T C G C T G A A A A G C T G T G G G A A A T C G G G |
| 13.92% | +1 | C A C A A A G T G G G A A A G A C C C A G C A T C \| n C G T G G G T C G C T G A A A A G C T G T G G G A A A T C G G |
| 7.33% | -5 | C A C A A A G T G G G A A A G A C C C A - - - - - \| C G T G G G T C G C T G A A A A G C T G T G G G A A A T C G G G |
| 6.2% | -8 | C A C A A A G T G G G A A A G A C C C A G C - - - \| - - - - - G T C G C T G A A A A G C T G T G G G A A A T C G G G |
| 5.19% | -4 | C A C A A A G T G G G A A A G A C C C A G C A T C \| - - - - G G T C G C T G A A A A G C T G T G G G A A A T C G G G |
| 3.55% | -6 | C A C A A A G T G G G A A A G A C C C A G C A T C \| - - - - - - T C G C T G A A A A G C T G T G G G A A A T C G G G |
| 3.42% | -2 | C A C A A A G T G G G A A A G A C C C A G C A T C \| - - T G G G T C G C T G A A A A G C T G T G G G A A A T C G G G |
| 3.15% | +13 | C A C A A A G T G G G A A A G A C C C A G C A T C \| n n n n n n n n n n n n n C G T G G G T C G C T G A A A A G C T |
| 2.94% | -5 | C A C A A A G T G G G A A A G A C C C A G C - - - \| - - T G G G T C G C T G A A A A G C T G T G G G A A A T C G G G |
| 2.81% | -20 | C A C A A A G T G G G - - - - - - - - - - - - - - \| - - - - - - T C G C T G A A A A G C T G T G G G A A A T C G G G |
| 2.58% | +15 | C A C A A A G T G G G A A A G A C C C A G C A T C \| n n n n n n n n n n n n n n n C G T G G G T C G C T G A A A A G |
| 2.56% | +10 | C A C A A A G T G G G A A A G A C C C A G C A T C \| n n n n n n n n n n C G T G G G T C G C T G A A A A G C T G T G |
| 1.78% | -4 | C A C A A A G T G G G A A A G A C C C A G - - - - \| C G T G G G T C G C T G A A A A G C T G T G G G A A A T C G G G |
| 1.76% | -13 | C A C A A A G T G G G A A A G A C C C A - - - - - \| - - - - - - - - G C T G A A A A G C T G T G G G A A A T C G G G |
| 1.72% | -7 | C A C A A A G T G G G A A A G A C C C A G C A T C \| - - - - - - - C G C T G A A A A G C T G T G G G A A A T C G G G |
| 1.72% | +9 | C A C A A A G T G G G A A A G A C C C A G C A T C \| n n n n n n n n n C G T G G G T C G C T G A A A A G C T G T G G |
| 1.37% | -3 | C A C A A A G T G G G A A A G A C C C A G C A T C \| - - - G G G T C G C T G A A A A G C T G T G G G A A A T C G G G |
| 1.21% | +8 | C A C A A A G T G G G A A A G A C C C A G C A T C \| n n n n n n n n C G T G G G T C G C T G A A A A G C T G T G G G |
| 0.85% | +14 | C A C A A A G T G G G A A A G A C C C A G C A T C \| n n n n n n n n n n n n n n C G T G G G T C G C T G A A A A G C |
| 0.74% | -13 | C A C A A A G T G G G A A A G A C C C A G C A T C \| - - - - - - - - - - - - - A A A G C T G T G G G A A A T C G G G |
| 0.62% | -6 | C A C A A A G T G G G A A A G A C C C A G C - - - \| - - - G G G T C G C T G A A A A G C T G T G G G A A A T C G G G |
| 0.5% | -2 | C A C A A A G T G G G A A A G A C C C A G C A - - \| C G T G G G T C G C T G A A A A G C T G T G G G A A A T C G G G |

**Supplementary figure 12 | Determination of editing efficiency in the HEKs1 locus by Inference of CRISPR Edits (ICE from SYNTHEGO) after transduction with VECTRv2-Cas(sgHEKs1).** An amplicon obtained from HEK293-EGFP cells transduced with a two-component VECTRv2-Cas(sgHEKs1) was analyzed by ICE. PCR product from mock-transduced cells was used as a control. Additional data to Fig. 3a (**a**) Summary of editing results. (**b**) Sanger sequencing chromatograms of the edited (upper panel) and control (lower panel) samples. The horizontal black underlined region represents the guide region. The PAM is underlined in red. The cut site is is shown by the vertical dotted line. (**c**) The left panel shows the level of disagreement between the control and edited samples around the cut site. Distribution of indel sizes and their frequencies is depicted in the right panel. (**d**) The contribution tab shows the sequences present in the pool of edited sequences. Their relative abundance is shown on the left. The cut site is represented by the red vertical dotted line.

#### SUPPLEMENTARY FIGURE 13

**a**

HEK293-EGFP cells

| Label | ICE (indel %) | KO-Score* | $R^2$ ** |
|---|---|---|---|
| LentiCRISPRv2(sgHEKs1) | 97 | 86 | 0.97 |

* Proportion of indels that indicate a frameshift or are 21+bp in length
** How well the proposed indel distribution fits the Sanger sequence data of the edited sample

**b**

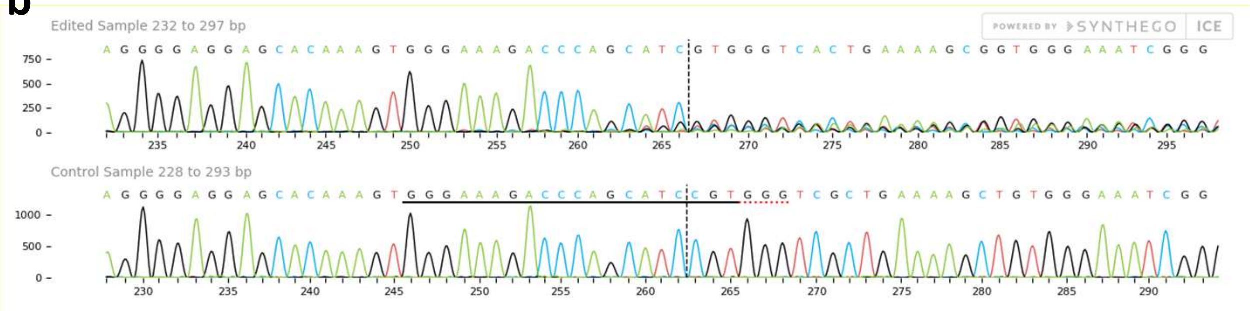


**c**

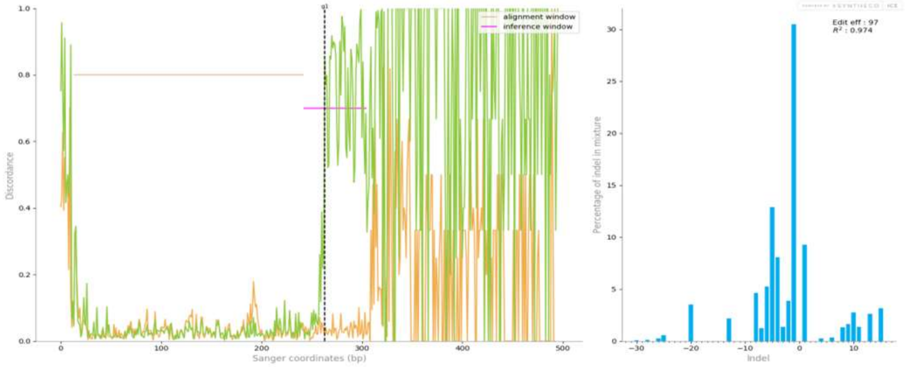


**d**

```
                       PAM
wt                     CACAAAGTGGGAAAGACCCAGCATC | CGTGGGTCGCTGAAAAGCTGTGGG

contribution  indel    sequence
30.5%         -1       CACAAAGTGGGAAAGACCCAGCAT- | CGTGGGTCGCTGAAAAGCTGTGGG
9.64%         -5       CACAAAGTGGGAAAGACCCA----- | CGTGGGTCGCTGAAAAGCTGTGGG
9.29%         +1       CACAAAGTGGGAAAGACCCAGCATC | nCGTGGGTCGCTGAAAAGCTGTGG
5.44%         -4       CACAAAGTGGGAAAGACCCAG---- | CGTGGGTCGCTGAAAAGCTGTGGG
5.24%         -6       CACAAAGTGGGAAAGACCCAGCATC | ------TCGCTGAAAAGCTGTGGG
4.63%         -8       CACAAAGTGGGAAAGACCCAGC--- | -----GTCGCTGAAAAGCTGTGGG
3.89%         -2       CACAAAGTGGGAAAGACCCAGCATC | --TGGGTCGCTGAAAAGCTGTGGG
3.25%         -5       CACAAAGTGGGAAAGACCCAGC--- | --TGGGTCGCTGAAAAGCTGTGGG
3.17%         + 15     CACAAAGTGGGAAAGACCCAGCATC | nnnnnnnnnnnnnnnCGTGGGTCG
2.75%         + 10     CACAAAGTGGGAAAGACCCAGCATC | nnnnnnnnnnCGTGGGTCGCTGAA
2.65%         -4       CACAAAGTGGGAAAGACCCAGCATC | ----GGTCGCTGAAAAGCTGTGGG
2.64%         +13      CACAAAGTGGGAAAGACCCAGCATC | nnnnnnnnnnnnnCGTGGGTCGCT
2.45%         -20      CACAAAGTGGG-------------- | ------TCGCTGAAAAGCTGTGGG
1.71%         -13      CACAAAGTGGGAAAGACCCA----- | --------GCTGAAAAGCTGTGGG
1.66%         + 9      CACAAAGTGGGAAAGACCCAGCATC | nnnnnnnnnCGTGGGTCGCTGAAA
1.36%         + 11     CACAAAGTGGGAAAGACCCAGCATC | nnnnnnnnnnnCGTGGGTCGCTGA
1.34%         + 8      CACAAAGTGGGAAAGACCCAGCATC | nnnnnnnnCGTGGGTCGCTGAAAA
1.26%         -7       CACAAAGTGGGAAAGACCCAGCATC | -------CGCTGAAAAGCTGTGGG
1.08%         -20      CACAAAGTGGGAAAGACCCA----- | ---------------AGCTGTGGG
1.01%         -3       CACAAAGTGGGAAAGACCCAGC--- | CGTGGGTCGCTGAAAAGCTGTGGG
```

**Supplementary figure 13 | Determination of editing efficiency in the HEKs1 locus by Inference of CRISPR Edits (ICE from SYNTHEGO) after transduction with LentiCRISPRv2(sgHEKs1).** An amplicon obtained from HEK293-EGFP cells transduced with a control LentiCRISPRv2(sgHEKs1) was analyzed by ICE. PCR product from mock-transduced cells was used as a control. Additional data to Fig. 3a (**a**) Summary of editing results. (**b**) Sanger sequencing chromatograms of the edited (upper panel) and control (lower panel) samples. The horizontal black underlined region represents the guide region. The PAM is underlined in red. The cut site is is shown by the vertical dotted line. (**c**) The left panel shows the level of disagreement between the control and edited samples around the cut site. Distribution of indel sizes and their frequencies is depicted in the right panel. (**d**) The contribution tab shows the sequences present in the pool of edited sequences. Their relative abundance is shown on the left. The cut site is represented by the red vertical dotted line.

#### SUPPLEMENTARY FIGURE 14

**a**

Jurkat cells

| Label | ICE (indel %) | KO-Score * | R2 ** |
|---|---|---|---|
| VECTRv2-Cas(sgFANCF) | 43 | 38 | 0.95 |

* Proportion of indels that indicate a frameshift or are 21+bp in length
** How well the proposed indel distribution fits the Sanger sequence data of the edited sample

**b**

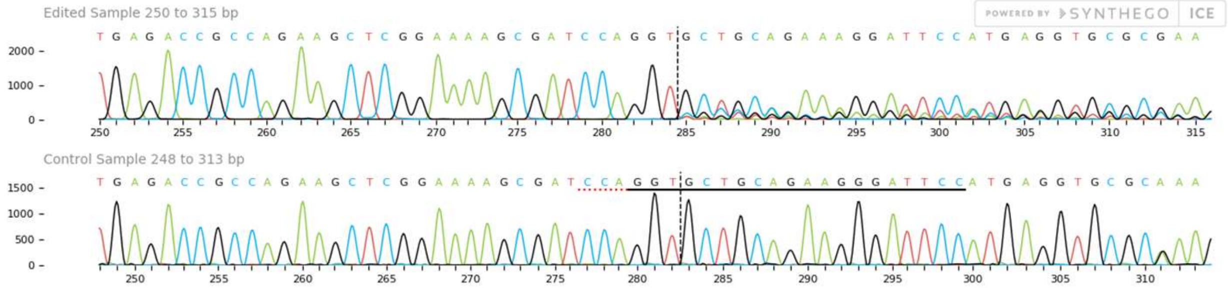


**c**

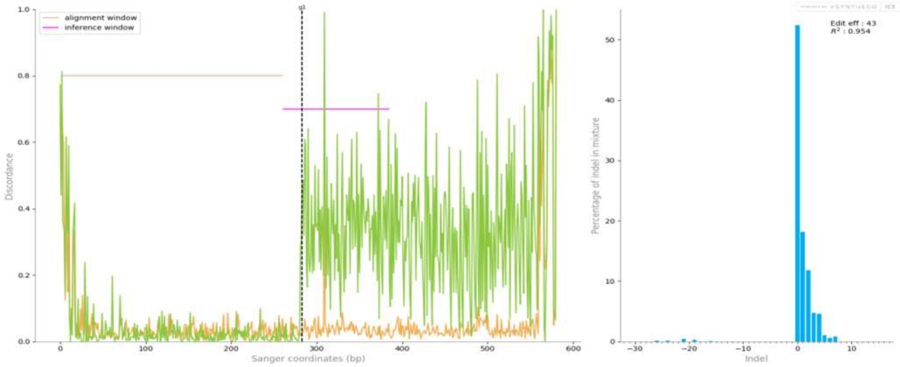


**d**

```
                              PAM
        wt    AGAAGCTCGGAAAAGCGATCCAGGT|GCTGCAGAAGGGATTCCATGAGGTGCG

52.43%   0    AGAAGCTCGGAAAAGCGATCCAGGT|GCTGCAGAAGGGATTCCATGAGGTGCG
18.17%   1    AGAAGCTCGGAAAAGCGATCCAGGT|nGCTGCAGAAGGGATTCCATGAGGTGC
11.82%   2    AGAAGCTCGGAAAAGCGATCCAGGT|nnGCTGCAGAAGGGATTCCATGAGGTG
4.73%    3    AGAAGCTCGGAAAAGCGATCCAGGT|nnnGCTGCAGAAGGGATTCCATGAGGT
4.58%    4    AGAAGCTCGGAAAAGCGATCCAGGT|nnnnGCTGCAGAAGGGATTCCATGAGG
1.08%    5    AGAAGCTCGGAAAAGCGATCCAGGT|nnnnnGCTGCAGAAGGGATTCCATGAG
0.84%    7    AGAAGCTCGGAAAAGCGATCCAGGT|nnnnnnnGCTGCAGAAGGGATTCCATG
0.58%    6    AGAAGCTCGGAAAAGCGATCCAGGT|nnnnnnGCTGCAGAAGGGATTCCATGA
0.43%  -21    AGAAGCTCGGAAAAGCGA-------|--------------TCCATGAGGTGCG
0.28%  -19    AGAAGCTCGGAAAAGCGATCC----|---------------CCATGAGGTGCG
0.17%  -24    AGAAGCTCGGAAAAGC---------|---------------CCATGAGGTGCG
0.13%  -26    AGAAGCTCGGAAAA-----------|---------------CCATGAGGTGCG
0.08%  -16    AGAAGCTCGGAAAAGCGATCCAG--|--------------TCCATGAGGTGCG
0.03%  -21    AGAAGCTCGGAAAAGC---------|------------ATTCCATGAGGTGCG
```

**Supplementary figure 14 | Determination of editing efficiency in the FANCF locus by Inference of CRISPR Edits (ICE from SYNTHEGO) after transduction with VECTRv2-Cas(sgFANCF).** An amplicon obtained from Jurkat cells transduced with a two-component VECTRv2-Cas(sgFANCF) was analyzed by ICE. PCR product from mock-transduced cells was used as a control. Additional data to Fig. 3b (**a**) Summary of editing results. (**b**) Sanger sequencing chromatograms of the edited (upper panel) and control (lower panel) samples. The horizontal black underlined region represents the guide region. The PAM is underlined in red. The cut site is is shown by the vertical dotted line. (**c**) The left panel shows the level of disagreement between the control and edited samples around the cut site. Distribution of indel sizes and their frequencies is depicted in the right panel. (**d**) The contribution tab shows the sequences present in the pool of edited sequences. Their relative abundance is shown on the left. The cut site is represented by the red vertical dotted line.

#### SUPPLEMENTARY FIGURE 15

**a**

Jurkat cells

| Label | ICE (indel %) | KO-Score* | $R^2$ ** |
|---|---|---|---|
| LentiCRISPRv2(sgFANCF) | 51 | 43 | 0.95 |

* Proportion of indels that indicate a frameshift or are 21+bp in length
** How well the proposed indel distribution fits the Sanger sequence data of the edited sample

**b**

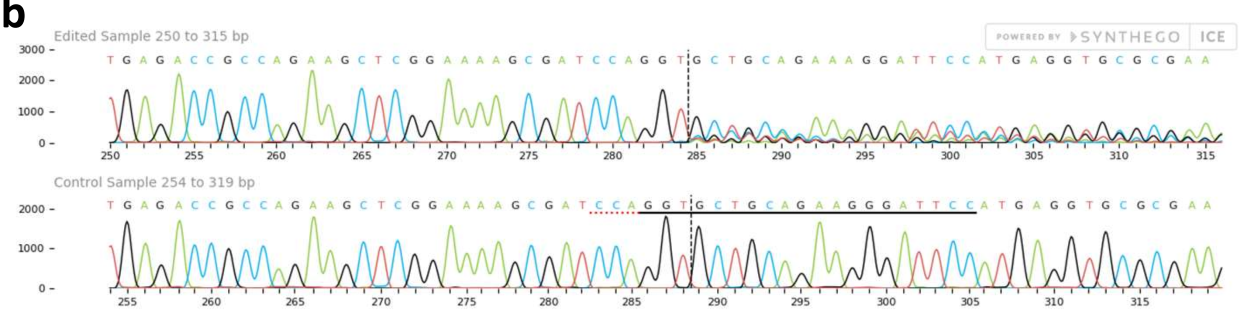


**c**

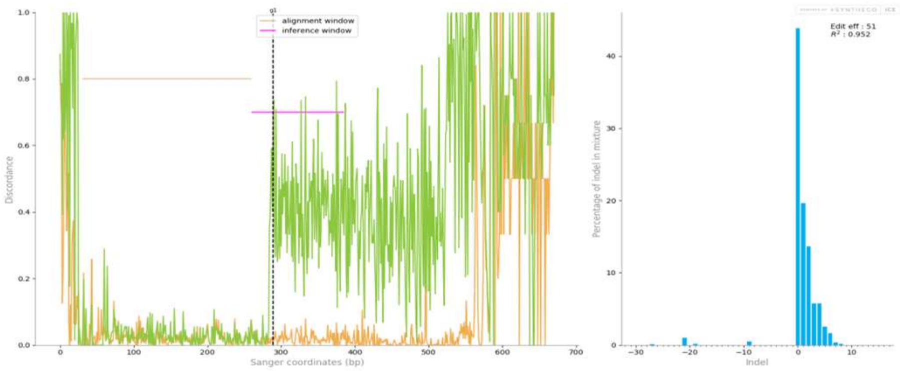


**d**

PAM

| | | |
|---|---|---|
| | wt | A G A A G C T C G G A A A A G C G A T C C A G G T \| G C T G C A G A A G G G A T T C C A T G A G G T G C G |
| 43.86% | 0 | A G A A G C T C G G A A A A G C G A T C C A G G T \| G C T G C A G A A G G G A T T C C A T G A G G T G C G |
| 19.62% | +1 | A G A A G C T C G G A A A A G C G A T C C A G G T \| n G C T G C A G A A G G G A T T C C A T G A G G T G C |
| 13.66% | +2 | A G A A G C T C G G A A A A G C G A T C C A G G T \| n n G C T G C A G A A G G G A T T C C A T G A G G T G |
| 5.76% | +3 | A G A A G C T C G G A A A A G C G A T C C A G G T \| n n n G C T G C A G A A G G G A T T C C A T G A G G T |
| 5.74% | +4 | A G A A G C T C G G A A A A G C G A T C C A G G T \| n n n n G C T G C A G A A G G G A T T C C A T G A G G |
| 2.58% | + 5 | A G A A G C T C G G A A A A G C G A T C C A G G T \| n n n n n G C T G C A G A A G G G A T T C C A T G A G |
| 1.68% | + 6 | A G A A G C T C G G A A A A G C G A T C C A G G T \| n n n n n n G C T G C A G A A G G G A T T C C A T G A |
| 0.83% | -21 | A G A A G C T C G G A A A A G C - - - - - - - - - \| - - - - - - - - - - - - A T T C C A T G A G G T G C G |
| 0.47% | -9 | A G A A G C T C G G A A A A G C G A T C - - - - - \| - - - - C A G A A G G G A T T C C A T G A G G T G C G |
| 0.39% | + 7 | A G A A G C T C G G A A A A G C G A T C C A G G T \| n n n n n n n G C T G C A G A A G G G A T T C C A T G |
| 0.2% | + 8 | A G A A G C T C G G A A A A G C G A T C C A G G T \| n n n n n n n n G C T G C A G A A G G G A T T C C A T |
| 0.16% | -21 | A G A A G C T C G G - - - - - - - - - - - - - - - \| - - - - - - G A A G G G A T T C C A T G A G G T G C G |
| 0.15% | -19 | A G A A G C T C G G A A A A G C G A T C C - - - - \| - - - - - - - - - - - - - - - C C A T G A G G T G C G |
| 0.11% | -27 | A G A A G C T C G G A A A - - - - - - - - - - - - \| - - - - - - - - - - - - - - - C C A T G A G G T G C G |
| 0.01% | -21 | A G A A G C T C G G A A A - - - - - - - - - - - - \| - - - - - - - - - G G G A T T C C A T G A G G T G C G |

**Supplementary figure 15 | Determination of editing efficiency in the FANCF locus by Inference of CRISPR Edits (ICE from SYNTHEGO) after transduction with LentiCRISPRv2(sgFANCF).** An amplicon obtained from Jurkat cells transduced with a control LentiCRISPRv2(sgFANCF) was analyzed by ICE. PCR product from mock-transduced cells was used as a control. Additional data to Fig. 3b (**a**) Summary of editing results. (**b**) Sanger sequencing chromatograms of the edited (upper panel) and control (lower panel) samples. The horizontal black underlined region represents the guide region. The PAM is underlined in red. The cut site is is shown by the vertical dotted line. (**c**) The left panel shows the level of disagreement between the control and edited samples around the cut site. Distribution of indel sizes and their frequencies is depicted in the right panel. (**d**) The contribution tab shows the sequences present in the pool of edited sequences. Their relative abundance is shown on the left. The cut site is represented by the red vertical dotted line.

#### SUPPLEMENTARY FIGURE 16

**a**

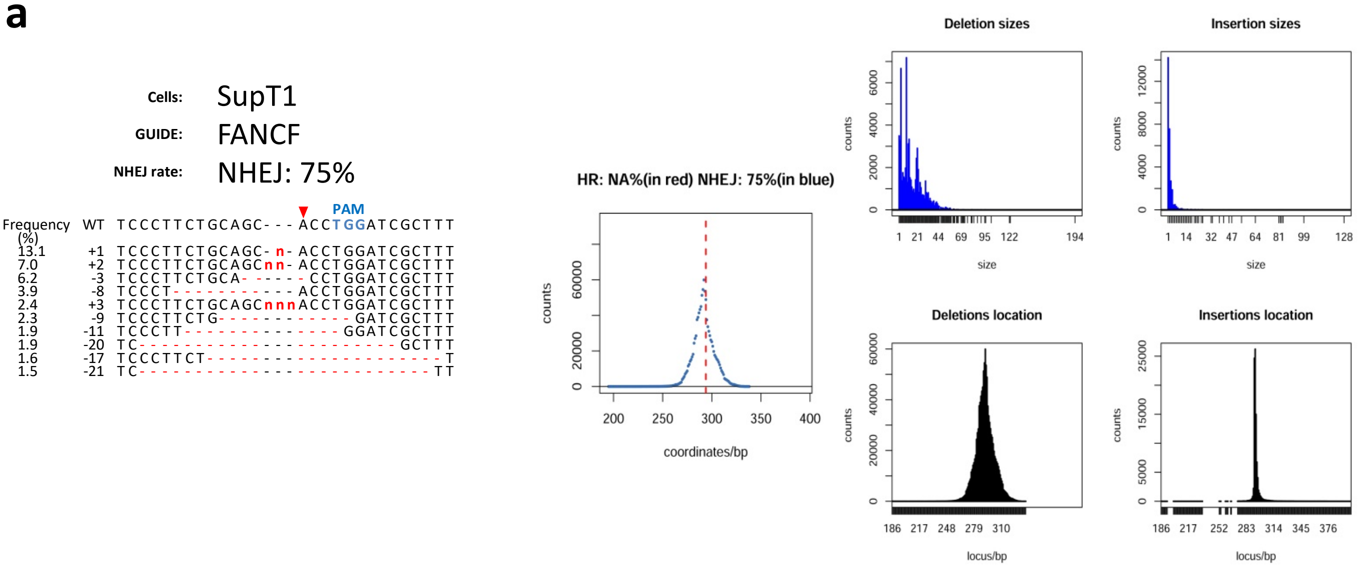


**b**

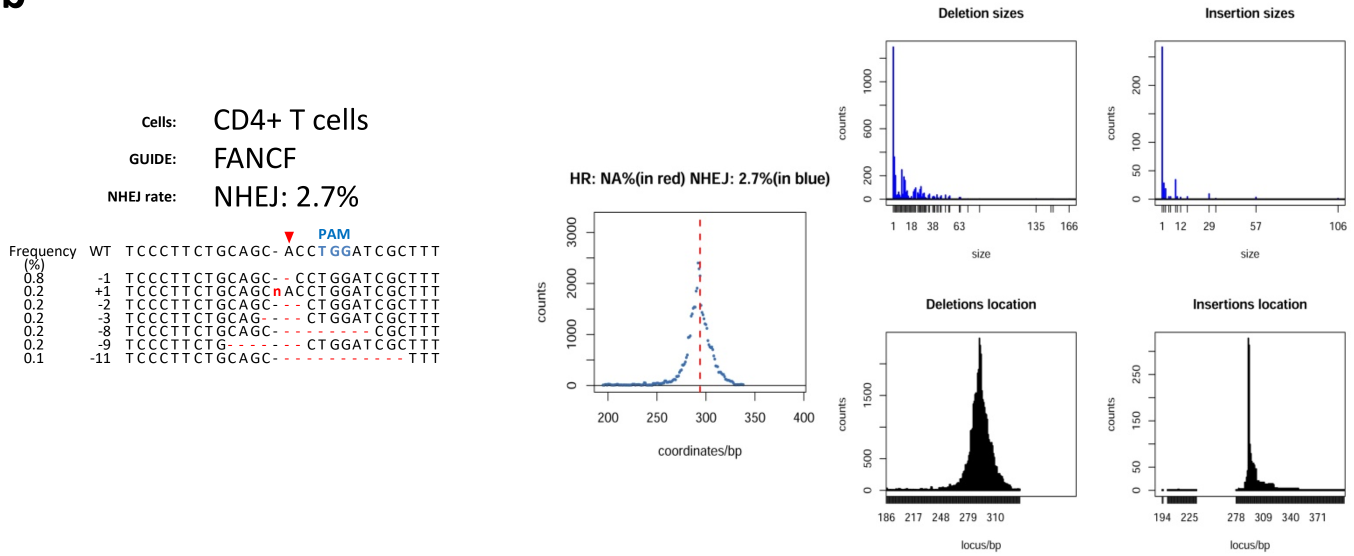


**Supplementary figure 16 | Frequency of VECTRv2-Cas(sgFANCF)-mediated indel formation by non-homologous end joining (NHEJ) in lymphocytes**. The rate of NHEJ at the double-strand breaks was assessed three days after transduction of SupT1 cell line (**a**) and primary CD4+ T cells (**b**) by high throughput sequencing combined with CRISPR-GA analysis (http://crispr-ga.net/) (Guell, Yang et al. 2014). Left panel: wild-type (WT) and mutated sequences at the FANCF locus. Deletions are indicated by the red dashes; insertions by "n"; PAM sequence is in blue. Frequency of occurrence of the sequence with insertion or deletion is shown at the left. Total indel frequency ("NEHJ:") is shown above the alignment. Middle panel: number of deletions and homologous recombination (HR) events at each corresponding location. HR and NHEJ efficiency and is shown above the plot. Upper right panels show the number of insertions and deletions and their corresponding size. Bottom right panels display the number of deletions and insertions and their corresponding location within the genomic locus of interest.

#### SUPPLEMENTARY FIGURE 17

**Cells:** SupT1

**GUIDE:** HEKs1

**NHEJ rate:** NHEJ: 100%

| Frequency (%) | | PAM |
|---|---|---|
| | WT | GAAAGACCCAGCATC--CGTGGGTCGCTGAAAAG |
| 14.5 | +1 | GAAAGACCCAGCATCn-CGTGGGTCGCTGAAAAG |
| 11.5 | -1 | GAAAGACCCAGCAT---CGTGGGTCGCTGAAAAG |
| 10 | +2 | GAAAGACCCAGCATCnnCGTGGGTCGCTGAAAAG |
| 5 | -13 | GAAAGACCCAGC---------------TGAAAAG |
| 3.5 | -8 | GAAAGACCCAGCAT----------CGCTGAAAAG |
| 3 | -4 | GAAAGACCCAGC------GTGGGTCGCTGAAAAG |
| 3 | -10 | GAAAGACCCAGCAT------------CTGAAAAG |
| 3 | -26 | GAAA----------------------------AG |
| 2.5 | -2 | GAAAGACCCAGCA----CGTGGGTCGCTGAAAAG |
| 2.5 | -9 | GAAAGAC-----------GTGGGTCGCTGAAAAG |
| 2.5 | -12 | GAAAGACCCAGCAT--------------GAAAAG |
| 2.5 | -17 | GAAAGACCG-------------------GAAAAG |
| 2.5 | -18 | GAAAGAC--------------------TGAAAAG |

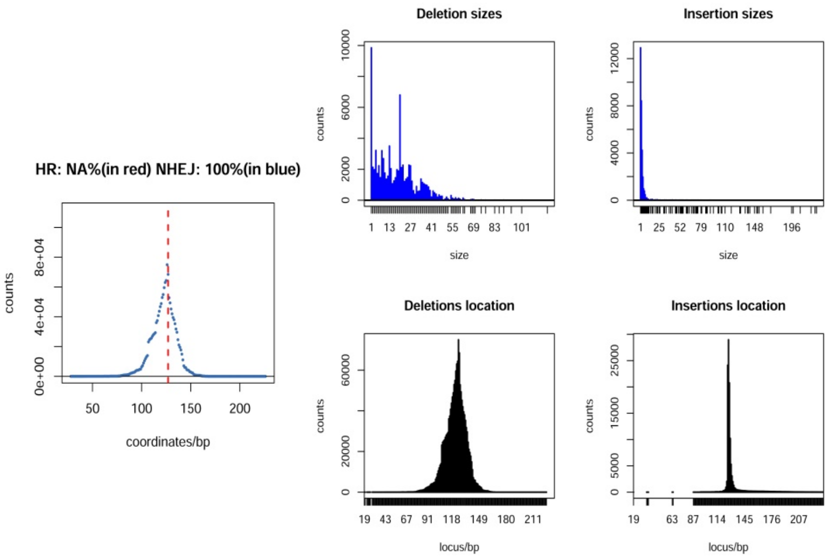


**Cells:** CD4+ T cells

**GUIDE:** HEKs1

**NHEJ rate:** NHEJ: 15%

| Frequency (%) | | PAM |
|---|---|---|
| | WT | GAAAGACCCAGCATC-CGTGGGTCGCTGAAAAG |
| 1.8 | -1 | GAAAGACCCAGCATC--GTGGGTCGCTGAAAAG |
| 1.4 | -20 | G---------------------TCGCTGAAAAG |
| 0.9 | +1 | GAAAGACCCAGCATCnCGTGGGTCGCTGAAAAG |
| 0.8 | -13 | GAAAGACCCAGC--------------TGAAAAG |
| 0.6 | -4 | GAAAGACCCAGC-----GTGGGTCGCTGAAAAG |
| 0.4 | -5 | GAAAGACCCAGC------TGGGTCGCTGAAAAG |
| 0.4 | -24 | GAAAGA-------------------------AG |
| 0.4 | -26 | GAAA----------------------------G |
| 0.3 | -27 | GAAAG---------------------------- |
| 0.3 | -10 | GAAAGACCCAGCATC-----------TGAAAAG |
| 0.3 | -2 | GAAAGACCCAGCA---CGTGGGTCGCTGAAAAG |
| 0.3 | -11 | GAAA------------CGTGGGTCGCTGAAAAG |
| 0.3 | -19 | G--------------------GTCGCTGAAAAG |

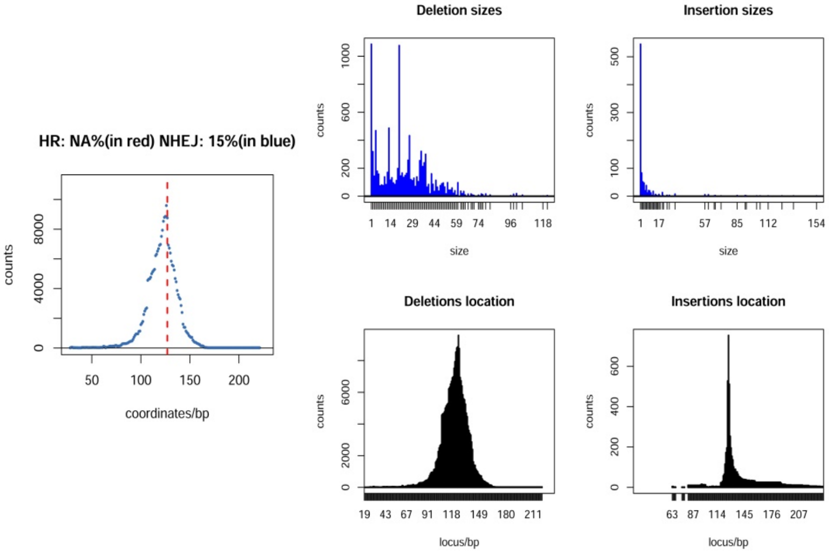


**Supplementary figure 17 | Frequency of VECTRv2-Cas(sgHEKs1)-mediated indel formation by non-homologous end joining (NHEJ) in lymphocytes**. The rate of NHEJ at the double-strand breaks was assessed three days after transduction of SupT1 cell line (**a**) and primary CD4+ T cells (**b**) by high throughput sequencing combined with CRISPR-GA analysis (http://crispr-ga.net/) (Guell, Yang et al. 2014). Left panel: wild-type (WT) and mutated sequences at the HEKs1 locus. Deletions are indicated by the red dashes; insertions by "n"; PAM sequence is in blue. Frequency of occurrence of the sequence with insertion or deletion is shown at the left. Total indel frequency ("NEHJ:") is shown above the alignment. Middle panel: number of deletions and homologous recombination (HR) events at each corresponding location. HR and NHEJ efficiency and is shown above the plot. Upper right panels show the number of insertions and deletions and their corresponding size. Bottom right panels display the number of deletions and insertions and their corresponding location within the genomic locus of interest.

#### SUPPLEMENTARY FIGURE 18

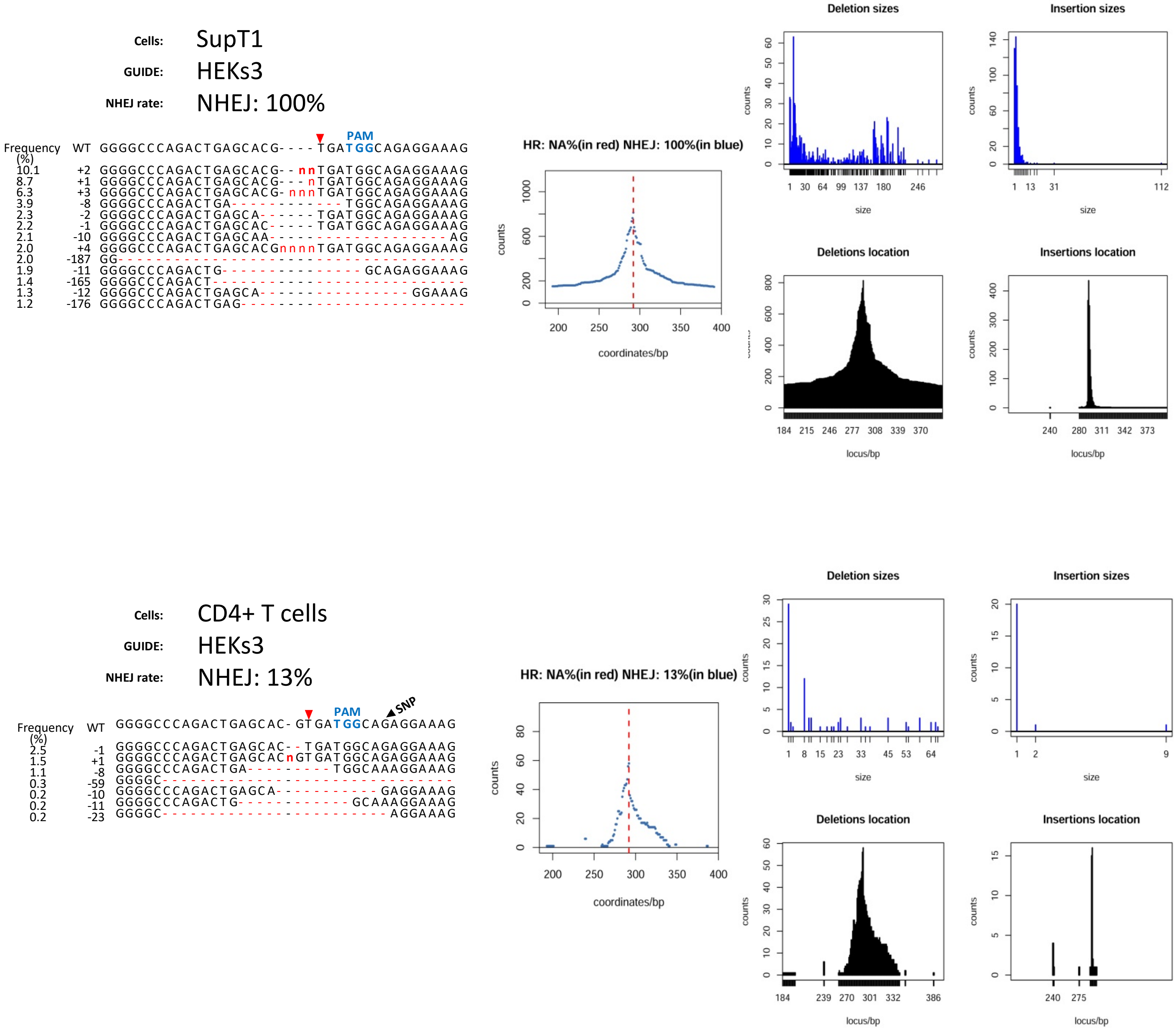


**Supplementary figure 18 | Frequency of VECTRv2-Cas(sgHEKs3)-mediated indel formation by non-homologous end joining (NHEJ) in lymphocytes.** The rate of NHEJ at the double-strand breaks was assessed three days after transduction of SupT1 cell line (**a**) and primary CD4+ T cells (**b**) by high throughput sequencing combined with CRISPR-GA analysis (http://crispr-ga.net/) (Guell, Yang et al. 2014). Left panel: wild-type (WT) and mutated sequences at the HEKs3 locus. Deletions are indicated by the red dashes; insertions by "n"; PAM sequence is in blue. Frequency of occurrence of the sequence with insertion or deletion is shown at the left. Total indel frequency ("NEHJ:") is shown above the alignment. Middle panel: number of deletions and homologous recombination (HR) events at each corresponding location. HR and NHEJ efficiency and is shown above the plot. Upper right panels show the number of insertions and deletions and their corresponding size. Bottom right panels display the number of deletions and insertions and their corresponding location within the genomic locus of interest.

### A novel toolkit for the efficient delivery of Cas9/sgRNA complexes to chromosomes in cells

#### MATERIAL AND METHODS

##### Cell lines, cell culture, and quantification of EGFP-expressing cells

HEK293T (ATCC CRL-3216), HEK293 cell (ATCC CRL-1573), IM9 (CCL-159), SupT1 (CRL-1942), Jurkat E6-1 (TIB-152), were obtained from ATCC. The THP-1 cells were obtained from Henning Hofmann (Robert Koch Institut). The human kidney cell lines were maintained in stable glutamine-containing high glucose Dulbecco's modified Eagle's medium (DMEM, Thermo Fisher Scientific) supplemented with 10 % fetal bovine serum (FBS Gold Plus, Bio-Sell). The cell lines derived from human lymphocytes and monocytes were cultivated in stable glutamine-containing RPMI-1640 (Carl Roth) supplemented with 10 % FBS Gold Plus (Bio-Sell). Cryopreserved Human CD4+ T cells from normal human peripheral blood were acquired from Zen-Bio. More than 95 % of the cells expressed CD3. The cells were cultured in X-VIVO 15 (Biozym) + 5 % FBS Gold Plus (Bio-Sell) supplemented with IL-2 (100 ng / ml; PEPROTech) and IL-7 (15 ng /ml; PEPROTech). The cells were activated one day prior to transduction by adding Dynabeads Human T-cell activator CD3/CD28 (Thermo Fisher Scientific) at a bead to cell ratio of 1:1. For all cultivated cells, no antibiotics were used. The cells were maintained at 37°C and 5 % $CO_2$ in a humidified incubator. To detect EGFP-positive cells, UV microscopy with the Olympus IX70 equipped with Olympus XM10 camera and CellSens software (Olympus). 947 ms acquisition time was used to detect the HEK293-EGFP cells expressing a low-level of EGFP. To quantify the number of EGFP-positive cells, flow cytometry was performed. The cells were trypsinized, washed and analyzed using FACSCalibour flow cytometer and CellQuest Pro Software (BD Biosciences). Forward versus side scatter gating was used to exclude debris and death cells from the analysis.

##### Plasmid construction

A pVpr.Prot.Cas9 plasmid was constructed by Gibson assembly of a gBlock ordered from IDT (containing Vpr, Protease cleavage site and SV40 nuclear localization signal–coding sequence) and two PCR products containing Cas9 coding sequence and Rev-responsive element (RRE), respectively. The pLentiCRISPRv2 (available from Addgene; plasmid #52961) was used as a template for the amplification of a DNA fragment encompassing the Cas9-coding sequence. The pCMgpRRE plasmid served as a template for the amplification of DNA fragment containing the RRE, CMV promoter, β globin polyA, and plasmid backbone sequences (Konstantoulas et.al., 2014). A Vpr-coding region from HIV-1 YU2 was used as a basis for the design of the gBlock sequence (Wu et al., 1995). A lentiviral transfer vector, Lenti(sgFILLER), which contains a filler sequence flanked by *Bsm*BI sites in place of the 20bp-long targeting crRNA sequence, was constructed from the pLentiCRISPRv2 by deleting Cas9-coding sequence in a long template PCR (primer pair: Fwd: 5'ATG ACC GAG TAC AAG CCC ACG3' ; Rev: 5' CCT GTG TTC TGG CGG CAA AC 3') followed by circularization of the resulting PCR product. Lentiviral transfer vectors carrying sgRNAs targeting specific loci in the genome (Lenti(sgRNA)) were constructed from the Lenti(sgFILLER). The vector was digested with *Bsm*BI and a pair of annealed and phosphorylated oligos was cloned into the single guide scaffold. D64V mutation was introduced into the psPAX2 plasmid by a high-fidelity PCR with a primer pair carrying the desired nucleotide substitution. Plasmids were amplified in DH5α or NEB Stable competent *E.coli* (New England Biolabs) and purified using a Qiagen Plasmid Midi kit (Qiagen).

**Lentivector production and transduction**

To produce lentivectors containing both Cas9 protein and U6-sgRNA template five plasmids (a total amount of 3.9 µg) were co-transfected to exponentially growing HEK293T cells seeded in 6-well plates (ATTC CRL-3216; ~80% confluent) using Fugene HD (Promega). We used 1.2 of transfer vector DNA (pLenti(sgRNA)), 0.8 µg of psPAX2 (Addgene #12260) or psPAX2D64V, 0.6 µg pRSV-Rev (Addgene #12253), 0.4 µg of pHCMV-G (Konstantoulas et al., 2014), and 0.9 µg of pVpr.Prot.Cas9. Transfection medium was replaced with fresh cell culture medium 20 h post transfection. Virus-containing supernatants were harvested 48 h post transfection, centrifuged (3500 rpm for 3 min), filtered (0.45 µm, Sarstedt), and immediately used. For transduction of adherent cells, ~5 x $10^4$ cells were plated in each well of a 12-well plate one day before transduction. The plated cells were incubated with the virus-containing supernatant (350 µl) supplemented with polybrene (8 µg/ml) for 6 h and fresh cell culture medium (DMEM + 10% FBS Gold Plus)) was added. For transduction of suspension cells, ~1 x $10^5$ cells in 50 µl of cell culture medium were incubated with 100 µl of virus-containing supernatant for 6h. Next, cells were pelleted and re-suspended in fresh cell culture medium.

**T7 endonuclease I assay**

To detect genomic modification at the targeted regions, genomic DNA was extracted from transduced cells 3 days post-transduction using the Quick Extract DNA Extraction Solution (Lucigen) and used for PCR to amplify specific on-target sites with Phusion high fidelity DNA polymerase (New England Biolabs) and primer pairs specified in **Supplementary information (list of primers)**. PCR products were purified by Ampure XP beads (Beckman Coulter) according to the manufacturer's instructions. 200 ng of purified DNA were denatured and hybridized (95˚C, 5min; 95̊C to 25̊C, -0.1̊C / s; hold at 4̊C) in 1x NEBuffer 2 (New England Biolabs) in a total volume of 14 µl. 1 µl of T7 Endonuclease I (New England Biolabs) was added to the hybridized PCR product and incubated at 37̊C for 30 min. 5 µl of 50 % glycerol was added to the T7 Endonuclease reaction and 20 µl was analyzed on a 2 % agarose gel containing peqGREEN (VWR). The DNA band intensity was quantified using VisionWorks LS Analysis Software. The frequency of indel formation was calculated using the following equation: $(1 – (1 – (b + c / a + b + c))^{1/2} ) \times 100$, where 'a' is the band intensity of DNA substrate and 'b'and 'c' are the cleavage products (Ran et al., 2013). It should be noted that we used this equation even though we are aware that it underestimates the editing efficiency in such a case that one type of editing predominates in a highly mutated locus as one mutant DNA duplex upon denaturation and re-annealing produces again mutant:mutant hybrid.

**High throughput sequencing and data analysis**

For deep sequencing, genomic DNA from SupT1 and CD4+ T cells was prepared as described above. The genomic region flanking the targeted site was amplified in 20 cycles using Phusion high fidelity DNA polymerase (New England Biolabs) and the primer pairs specified in **Supplementary**

**information (list of oligos)**. The amplified sequences were purified (Ampure XP beads (Beckman Coulter)) and send for library preparation and sequencing on a MiSeq high-throughput sequencer (2 x 300 bp; Illumina) to LGC Genomics (Berlin). The 300 bp paired-end MiSeq raw reads were de-multiplexed and low quality reads (a PHRED quality score of less than 30) removed using NextGen Sequence Workbench (Avalanche NextGen). The R1 and R2 fastq files were then uploaded to CRISPR Genome Analyzer (http://crispr-ga.net/) (Guell et al., 2014) together with the target sequence. The output figures and aligned indel-containing sequences flanking the cut site are shown in **Supplementary Figures 15 – 17**. Deep sequencing data is available at the NCBI's Sequencing Read Archive (SRA).

#### LIST OF OLIGOS

Primers used for the preparation of amplicons for high throughput sequencing:

| Name | oligo |
|---|---|
| HEK293s1F600 | 5' CTTGTCGGCAGTAGTGGGAG 3‘ |
| HEK293s1_R | 5' ACCAGCGTCTTCCCTTCCTC 3‘ |
| FANCFon600_R | 5' ATCATCTCGCACGTGGTTCC 3‘ |
| FANCF_F | 5' CCAGGTGCTGACGTAGGTAG 3‘ |
| HEK293s3F600 | 5' GGGTCACAGTGGCAAATGAG 3‘ |
| HEK293s3R600 | 5' TGTTGAGCTCGACCCTGAAG 3‘ |

Oligos used for the amplification of the targeted region in the *egfp* gene

| Name | oligo |
|---|---|
| CMV_sgRNA_F | ATCAATGGGCGTGGATAGCG |
| eGFP_sgRNA_R | TTGCCGTCCTCCTTGAAGTC |
| eGFP_sgRNA_R2 | GTCCATGCCGAGAGTGATCC |

Oligos used for cloning of target sequence into the *BsmBI*-digested Lenti(sgFILLER) backbone

| Name | oligo |
|---|---|
| EMX1sense | 5’caccgGAGTCCGAGCAGAAGAAGAA 3’ |
| EMX1anti | 5’aaaCTTCTTCTTCTGCTCGGACTCc 3’ |
| FANCFsense | 5’ caccgGGAATCCCTTCTGCAGCACC 3’ |
| FANCFanti | 5’aaaCGGTGCTGCAGAAGGGATTCCc 3’ |
| HEKs1sense | 5’ caccgGGGAAAGACCCAGCATCCGT 3’ |
| HEKs1anti | 5’aaaCACGGATGCTGGGTCTTTCCCc 3’ |
| HEKs3sense | 5’ caccgGGCCCAGACTGAGCACGTGA 3’ |
| HEKs3anti | 5’ aaaCTCACGTGCTCAGTCTGGGCCc 3’ |

#### LIST OF OLIGOS

Oligos used for cloning of target sequence into the *BsmBI*-digested Lenti(sgFILLER) backbone

| Name | oligo |
|---|---|
| EGFPsite2sense | 5' caccgGTCGCCCTCGAACTTCACCT 3' |
| EGFPsite3sense | 5' caccgGGCGAGGGCGATGCCACCTA 3' |
| EGFPsite4sense | 5' caccgGGTCGCCACCATGGTGAGCA 3' |
| EGFPsite5sense | 5' caccgGGTCAGGGTGGTCACGAGGG 3' |
| EGFPsite6sense | 5' caccgGGTGGTGCAGATGAACTTCA 3' |
| EGFPsite7sense | 5' caccgGTTGGGGTCTTTGCTCAGGG 3' |
| EGFPsite8sense | 5' caccgGATGCCGTTCTTCTGCTTGT 3' |
| EGFPsite2anti | 5' aaacAGGTGAAGTTCGAGGGCGACc 3' |
| EGFPsite3anti | 5' aaacTAGGTGGCATCGCCCTCGCCc 3' |
| EGFPsite4anti | 5' aaacTGCTCACCATGGTGGCGACCc 3' |
| EGFPsite5anti | 5' aaacCCCTCGTGACCACCCTGACCc 3' |
| EGFPsite6anti | 5' aaacTGAAGTTCATCTGCACCACCc 3' |
| EGFPsite7anti | 5' aaacCCCTGAGCAAAGACCCCAACc 3' |
| EGFPsite8anti | 5' aaacACAAGCAGAAGAACGGCATCc 3' |
| EGFPsite1_sense | 5' caccgGGGCGAGGAGCTGTTCACCG 3' |
| EGFPsite1_anti | 5' aaacCGGTGAACAGCTCCTCGCCCc 3' |
| EGFPsite1_mismatch_1&2_sense | 5' caccgGGGCGAGGAGCTGTTCACgc 3' |
| EGFPsite1_mismatch_3&4_sense | 5' caccgGGGCGAGGAGCTGTTCtgCG 3' |
| EGFPsite1_mismatch_5&6_sense | 5' caccgGGGCGAGGAGCTGTagACCG 3' |
| EGFPsite1_mismatch_7&8_sense | 5' caccgGGGCGAGGAGCTcaTCACCG 3' |
| EGFPsite1_mismatch_9&10_sense | 5' caccgGGGCGAGGAGgaGTTCACCG 3' |
| EGFPsite1_mismatch_11&12_sense | 5' caccgGGGCGAGGtcCTGTTCACCG 3' |
| EGFPsite1_mismatch_13&14_sense | 5' caccgGGGCGAccAGCTGTTCACCG 3' |
| EGFPsite1_mismatch_15&16_sense | 5' caccgGGGCctGGAGCTGTTCACCG 3' |
| EGFPsite1_mismatch_17&18_sense | 5' caccgGGcgGAGGAGCTGTTCACCG 3' |
| EGFPsite1_mismatch_18&19_sense | 5' caccgGccCGAGGAGCTGTTCACCG 3' |
| EGFPsite1_mismatch_1&2_anti | 5' aaacgcGTGAACAGCTCCTCGCCCc 3' |
| EGFPsite1_mismatch_3&4_anti | 5' aaacCGcaGAACAGCTCCTCGCCCc 3' |
| EGFPsite1_mismatch_5&6_anti | 5' aaacCGGTctACAGCTCCTCGCCCc 3' |
| EGFPsite1_mismatch_7&8_anti | 5' aaacCGGTGAtgAGCTCCTCGCCCc 3' |
| EGFPsite1_mismatch_9&10_anti | 5' aaacCGGTGAACtcCTCCTCGCCCc 3' |
| EGFPsite1_mismatch_11&12_anti | 5' aaacCGGTGAACAGgaCCTCGCCCc 3' |
| EGFPsite1_mismatch_13&14_anti | 5' aaacCGGTGAACAGCTggTCGCCCc 3' |
| EGFPsite1_mismatch_15&16_anti | 5' aaacCGGTGAACAGCTCCagGCCCc 3' |
| EGFPsite1_mismatch_17&18_anti | 5' aaacCGGTGAACAGCTCCTCcgCCc 3' |
| EGFPsite1_mismatch_18&19_anti | 5' aaacCGGTGAACAGCTCCTCGggCc 3' |
